# SARS-CoV-2 PLpro Hijacks a Conserved Stress Signaling Network to Drive Organ-Specific Loss of Epithelial Homeostasis

**DOI:** 10.64898/2026.03.24.713966

**Authors:** Quazi Taushif Ahmad, Shweta Banerjee, Isha Anerao, Saurabh Singh Parihar, Anjali Bajpai, Jyoti Tripathi, Mohd S Rizvi, Pradip Sinha

**Affiliations:** Biological Sciences and Bioengineering, Indian Institute of Technology, Kanpur; Department of Biomedical Engineering, Indian Institute of Technology, Hyderabad

**Keywords:** SARS-CoV-2, Papain-like protease, Drosophila, MDCK, epithelial barrier

## Abstract

Severe SARS-CoV-2 infection manifests as a systemic disorder characterized by catastrophic loss of epithelial homeostasis and dysregulated host inflammation. However, which specific viral factors initiate this tissue damage in intact, living organisms remains unresolved. Using a comprehensive *in vivo* screen of all 25 SARS-CoV-2 nonstructural proteins (NSPs) and accessory factors (ORFs) in the *Drosophila* wing imaginal epithelium, here we have identified NSP5 (Main protease, Mpro) and NSP3—specifically its papain-like protease (PLpro) domain—as individual triggers of tissue injury. Focusing further on PLpro, we demonstrate that its expression initiates a self-amplifying host cell stress signaling circuit driven by metabolic overdrive, oxidative stress, and hyperactivation of Akt, JNK, and JAK-STAT signaling. However, this pathogenic loop is fragile; genetic suppression of any of the hyperactivated signaling nodes collapses this network, thereby restoring epithelial homeostasis. This organ-intrinsic, PLpro-induced epithelial vulnerability is also seen in the adult midgut and larval respiratory system. Notably, in the larval respiratory system, PLpro drives compartment-specific pathologies: it induces fibrosis-like remodeling in terminally differentiated squamous tracheal tubes, while catastrophically depleting the progenitor-competent tracheoblasts and air sac precursors (ASP). Both these defects are reversed by the genetic knockdown of PLpro-induced JAK-STAT signaling. Finally, we show that PLpro expression in mammalian MDCK epithelia similarly triggers junctional remodeling and inflammatory stress signaling. Our findings position PLpro as a central driver of COVID-19 pathogenesis through a self-amplifying epithelial stress circuit that compromises epithelial homeostasis across organs and species.

## INTRODUCTION

SARS-CoV-2 infection disrupts epithelial homeostasis and often progress to multisystem disease (Lamers and Haagmans 2022; Berlin et al. 2020). In severe COVID-19, immune signaling becomes dysregulated, with impaired interferon responses and excessive inflammation, leading to multiorgan dysfunction (Blanco-Melo et al. 2020). A key feature of this pathology is loss of epithelial barrier integrity, along with disruption of tissue homeostasis and maladaptive inflammation (Deinhardt-Emmer et al. 2021; Gao et al. 2026; Hashimoto et al. 2022). Defining the molecular triggers of epithelial homeostasis failure is therefore critical to understanding COVID-19 pathogenesis.

A reductionist approach, testing one SARS-CoV-2 protein at a time in a host organ, can help identify which viral factors matter and which host pathways they engage. Although cell-based studies and interaction maps have identified many virus-host interactions (Gordon, Jang, et al. 2020; Gordon, Hiatt, et al. 2020; Stukalov et al. 2021; Hayn et al. 2021; Miao et al. 2021), they do not readily explain how individual viral proteins drive damage in intact tissues. As a result, these interactome maps often fail to connect viral protein function with loss of epithelial homeostasis or inflammatory injury *in vivo*.

To uncover mechanisms of pathogenesis—from cell-autonomous effects to systemic disease—requires an *in vivo* model that allows viral factors to be tested in intact tissue. *Drosophila* provides one such model. Many human proteins that interact with SARS-CoV-2 have conserved counterparts in *Drosophila* (van de Leemput and Han 2021; Guichard et al. 2023). This conservation underlies the *Drosophila* COVID-19 Resource (DCR), a transgenic toolkit for tissue-specific expression of individual viral factors in intact epithelia (Guichard et al. 2023). This approach allows the function of viral proteins to be studied outside the complexity of full viral replication and helps identify direct drivers of tissue damage.

Guided by this logic, we screened the fallouts of all twenty-five SARS-CoV-2 nonstructural proteins (NSPs) and accessory factors (ORFs) in the larval wing imaginal disc epithelium. This tissue is a useful model for epithelial homeostasis and cellular signaling crosstalks (Tripathi and Irvine 2022). Using cell death as the screening readout (Bergantiños et al. 2010; Goyal et al. 2000), we identified the papain-like protease (PLpro) domain of NSP3 as a major driver of epithelial tissue injury. We then tested PLpro in two other developmentally distinct organs, the larval respiratory system and the adult midgut. In both tissues, PLpro alone was sufficient to trigger a self-amplifying ROS-JNK-JAK-STAT injury circuit, in which metabolic strain, oxidative stress, and tissue-remodeling signals reinforced one another (Kucinski et al. 2017; Gain et al. 2023; Allen et al. 2022). Importantly, this circuit was fragile: blocking even a single host signaling pathway from this circuitry was sufficient to interrupt the injury loop, restore epithelial homeostasis, and preserve tissue architecture.

Further, in the respiratory epithelium, PLpro produced distinct effects in differentiated and progenitor-like cells. For instance, it drives fibrosis-like remodeling in differentiated dorsal tracheal tubes, while depleting the progenitor-competent tracheoblasts and air sac precursors (ASPs). This tissue-level vulnerability was also conserved in mammalian MDCK epithelial cells, where PLpro induced comparable stress and junctional remodeling.

Our findings identify SARS-CoV-2 PLpro as a disruptor of epithelial homeostasis via a self-amplifying host-stress signaling network. Beyond its established roles in viral replication and immune evasion, PLpro is sufficient to initiate tissue injury across distinct epithelial contexts *in vivo*. These results suggest that PLpro-induced pathology can be attenuated by suppressing individual nodes of the stress circuit, including JAK/STAT.

## RESULTS

### SARS-CoV-2 PLpro is sufficient to trigger injury-linked epithelial cell death and repair-like remodeling

Cleavage of the SARS-CoV-2 polyproteins pp1a and pp1ab by the papain-like protease (PLpro), encoded by NSP3, and the main protease, encoded by NSP5, generates 16 nonstructural proteins (NSPs) and 9 open reading frames (ORFs) (**Supplementary Figure 1A, A**□) (Osipiuk et al. 2021; Yadav et al. 2022). In the *Drosophila* COVID Resource (DRC), each transgenic NSP/ORF line can be conditionally expressed via the Gal4/UAS system, enabling *in vivo* analysis of functional interactions between individual viral proteins and host pathways (Guichard et al. 2023).

Because host–virus interactions frequently culminate in epithelial damage and cell death (Zheng et al. 2020; Lamers and Haagmans 2022), we used the wing imaginal disc as a sensitized *in vivo* platform to identify pathogenic SARS-CoV-2 proteins. Individual NSPs/ORFs were expressed along the dorsal–ventral boundary of the larval wing imaginal disc epithelium using *vg-Gal4* **(Supplementary Figure 2A, B**). Adult wings were then scored for the cut-wing phenotype, a classical readout of epithelial cell death in this tissue and a phenotype readily compared with that induced by Reaper (Bergantiños et al. 2010; Goyal et al. 2000) (**Supplementary Figure 2C**). Of the 25 NSPs/ORFs tested, only NSP3 and NSP5 produced the expected cut-wing phenotype (**Supplementary Figure 2E, F**).

Because NSP5 is already well characterized through its main protease function (Reinke et al. 2024; Biernacki et al. 2024; Bege and Borbás 2024), we focused subsequent analysis on NSP3. NSP3 expression in the wing imaginal disc epithelium, the D/V boundary displayed, as anticipated, activation of the effector caspase DrICE, visualized via immunoreactivity to cleaved Caspase-3 (Fraser et al. 1997) (**Supplementary Figure 2G**), reminiscent of caspase activation seen in SARS-CoV-2–infected human epithelium (Liang et al. 2024).

NSP3 contains a centrally located ∼35 kDa papain-like protease (PLpro) domain, which provides cysteine protease, deubiquitinating (DUB), and deISGylating activities (**Supplementary Figure 1B, B**□). PLpro is one of the most pathogenically important SARS-CoV-2 proteins: it processes pp1a to release proteins required for the replication–transcription complex (RTC), and it also dampens host innate immune responses, including type I interferon production (Shin et al. 2020; Cao et al. 2023; Liu et al. 2021). The remaining regions of NSP3, including Ubl1, the macrodomain, SUD, NAB, and the ectodomain (**Supplementary Figure 1B)**, are structurally distinct modules with other functions (Yang and Rao 2021; Shin et al. 2020). Because full-length NSP3 is a massive, multi-domain protein whose diverse structural modules could mask or confound domain-specific mechanisms or engage in non-proteolytic interactions, we used a targeted, domain-specific approach to pinpoint the driver of tissue injury. We thus generated a transgenic line expressing PLpro (see **SI**). Expression of *vg>PLpro* in the adult wing reproduced the cut-wing phenotype, localizing the cell death-inducing activity of NSP3 to its PLpro domain (**Figure 1A**). Co-expression of PLpro with Dronc^DN^, a dominant-negative form of the initiator caspase Dronc that inhibits apoptosome-mediated cell death (Shapiro et al. 2008), suppressed the cut-wing phenotype (**Figure 1A**), thereby linking PLpro-induced cell death to loss of adult wing pattern.

**Figure 1.**
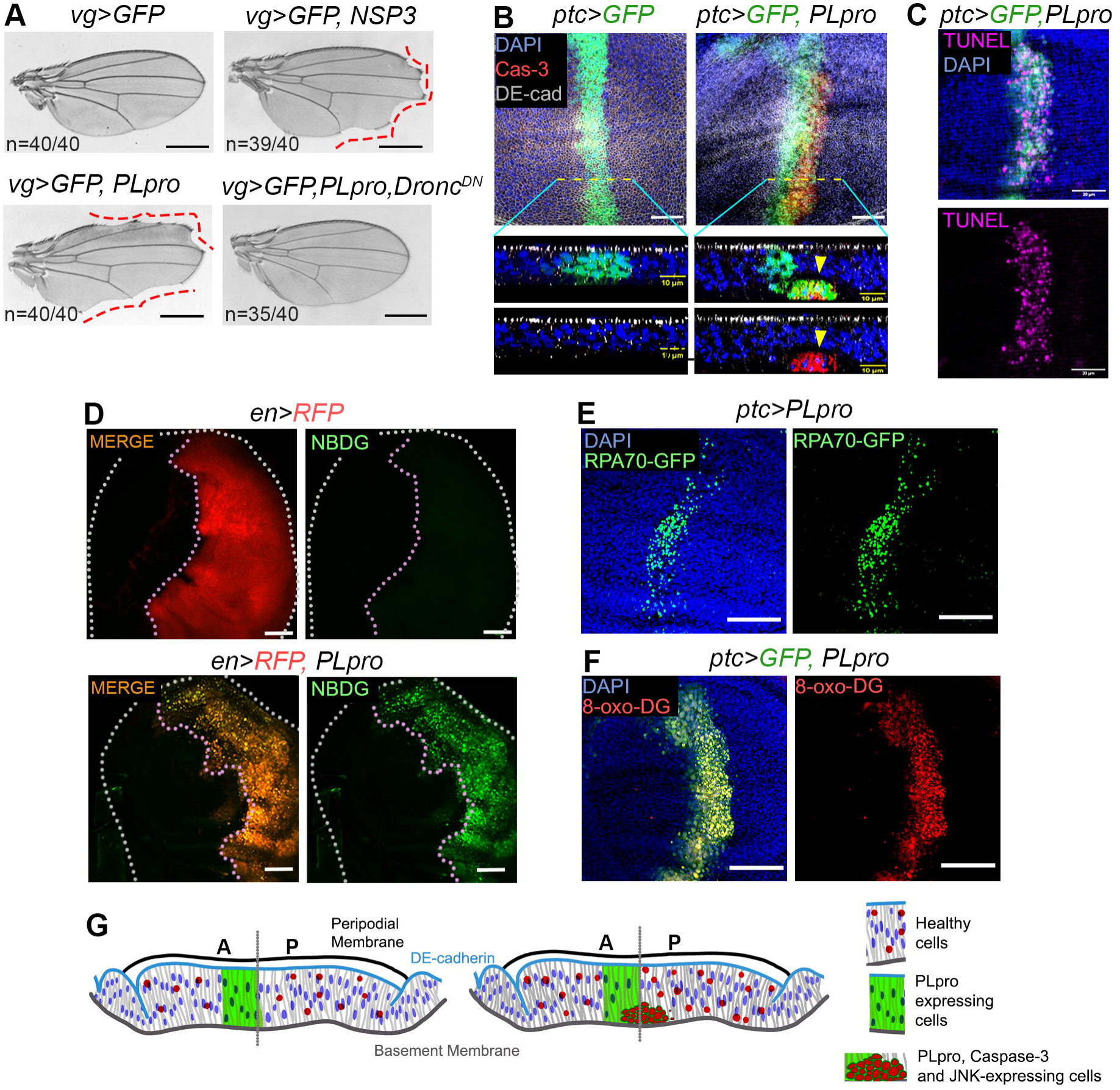
SARS-CoV-2 PLpro expression induces epithelial cell death and metabolic stress in the developing *Drosophila* wing epithelium. **(A)** Adult wings from *vg>GFP* (control), *vg>GFP, NSP3*, *vg>GFP, PLpro*, and *vg>GFP, PLpro, Dronc^DN^* flies. Dashed lines indicate wing margins displaying the cut-wing phenotype. **(B)** Confocal 3D projections of wing imaginal discs from control (*ptc>GFP*) and *ptc>GFP, PLpro* larvae stained for cleaved Caspase-3 (red). Corresponding X–Z optical sections along the indicated dashed lines are shown below each image. **(C)** Wing imaginal disc from *ptc>GFP, PLpro* larvae displaying TUNEL staining (magenta). **(D)** Wing imaginal discs from control (*en > RFP*) and *en > RFP, PLpro* larvae incubated with the fluorescent glucose analogue 2-NBDG (green) to assess glucose uptake. **(E)** Wing imaginal disc from *ptc>GFP, PLpro* larvae expressing the RPA70-GFP reporter (green), indicating replication stress-associated single-stranded DNA damage. **(F)** Wing imaginal disc from *ptc>GFP, PLpro* larvae stained for 8-oxo-dG (red), indicating oxidative DNA damage. **(G)** Schematic illustration of PLpro-induced epithelial remodeling in the wing imaginal disc. In control epithelia, columnar cells maintain apical–basal polarity, intact DE-cadherin-mediated cell-cell junctions, and attachment to the basement membrane. PLpro expression promotes epithelial disorganization, JNK activation, Caspase-3 cleavage, loss of cell-matrix adhesion, and basal cell extrusion. Elevated mitotic activity in the PLpro-expressing epithelium is indicated by an increased abundance of phospho-histone H3 (PH3)-positive cells (represented as red dots). Scale bars: 0.5 mm (A), 25 µm (B, D–F), 20 µm (C), and 10 µm (B, X–Z sections). n = 25 larvae per genotype (B–F). DAPI (blue) marks nuclei.

To trace the fate of PLpro-expressing cells in the developing epithelium, we examined *ptc>PLpro* wing imaginal discs **(Supplementary Figure 2D)**, focusing on the relatively flat wing pouch region along the anterior–posterior boundary where *ptc-Gal4* is active (**Supplementary Figure 2A, B**). In this pseudostratified epithelium, X–Z optical sections revealed three features along the apicobasal axis: PLpro-expressing cells (GFP+) within the main epithelial layer, GFP+ cells with caspase activation undergoing early basal delamination, and fully basal cells that retained cleaved caspase-3 staining but lacked GFP (**Figure 1B**). These observations capture a continuum in which PLpro-expressing epithelial cells activate caspase-3, delaminate basally, and are ultimately extruded. TUNEL labeling confirmed cell death in these PLpro-expressing cells (**Figure 1C**). Further, consistent with the coupled occurrence of cell death and regeneration in imaginal epithelia (Bergantiños et al. 2010; Klemm et al. 2021), the *ptc>PLpro* wing pouch showed non-autonomous induction of cell proliferation, although the normal pattern was not restored (**Supplementary Figure 3**). PLpro-induced epithelial cell death could reflect broader perturbations in host homeostasis, including metabolic stress and signaling rewiring that accompany epithelial injury and viral infection (Mullen et al. 2021; Codo et al. 2020; Lamers and Haagmans 2022). To examine this possibility, we assessed PLpro-expressing wing epithelium for altered metabolism, oxidative stress, and stress-activated signaling. PLpro expression was associated with increased glucose uptake (**Figure 1D**) (Codo et al. 2020), upregulation of the DNA repair marker RPA70-GFP (**Figure 1E**), and accumulation of 8-oxo-dG (Valavanidis et al. 2009), consistent with oxidative DNA damage and activation of downstream stress responses (Tian et al. 2021; Mullen et al. 2021) (**Figure 1F**).

These findings identify SARS-CoV-2 PLpro as a sufficient epithelial injury factor that acts independently of the full viral infection program to trigger cell death, delamination, and repair-like tissue remodeling *in vivo* (**Figure 1G**). A single SARS-CoV-2 domain thus reveals a conserved core injury-response program and uncovers a previously unrecognized pathogenic activity of PLpro.

### PLpro uncovers epithelial metabolic and stress-signaling reprogramming

Metabolic overdrive and genomic strain observed in the PLpro-expressing epithelium suggested an underlying surge in oxidative stress. To test this, we monitored reactive oxygen species (ROS) production. We found that PLpro-expressing epithelia displayed robust oxidative stress, as revealed by DHE (Fogarty et al. 2016) staining (**Figure 2A**), while knockdown of *Duox*, which encodes the ROS-producing enzyme Dual Oxidase (Fogarty et al. 2016), reduced caspase activity, and suppressed the adult cut-wing phenotype (**Figure 2B, C**).

**Figure 2.**
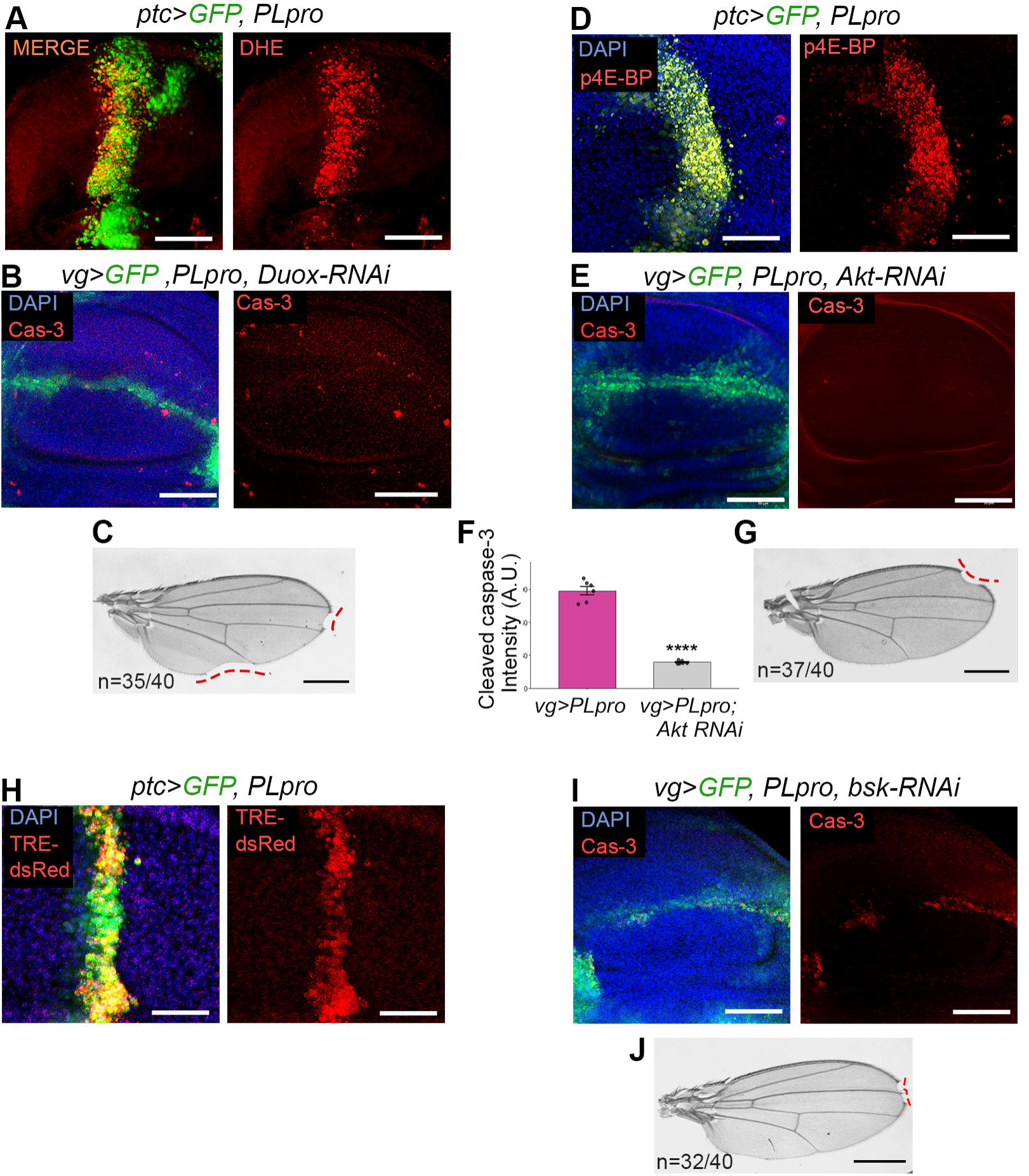
PLpro induces oxidative stress, Akt and JNK signaling, and epithelial cell death in the *Drosophila* wing disc. **(A)** Wing imaginal disc from *ptc>GFP, PLpro* larvae stained with dihydroethidium (DHE; red). **(B)** Wing imaginal disc from *vg>GFP, PLpro, Duox-RNAi* larvae immunostained for cleaved Caspase-3 (red). **(C)** Adult wing from *vg>GFP, PLpro, Duox-RNAi* flies. Dashed lines indicate wing margins displaying the cut-wing phenotype. **(D)** Wing imaginal disc from *ptc>GFP, PLpro* larvae stained for phosphorylated 4E-BP (p4E-BP; red). **(E)** Wing imaginal disc from *vg>GFP, PLpro, Akt-RNAi* larvae immunostained for cleaved Caspase-3 (red). **(F)** Quantification of cleaved Caspase-3 fluorescence intensity in *vg>GFP, PLpro* wing imaginal discs with or without *Akt* knockdown. Data represent mean ± s.d.; ****P<0.0001 by two-tailed unpaired Student’s *t*-test. **(G)** Adult wing from *vg>GFP, PLpro, Akt-RNAi* flies. Dashed lines indicate wing margins displaying the cut-wing phenotype. **(H)** Wing imaginal disc from *ptc>GFP, PLpro* larvae carrying the TRE-dsRed reporter. **(I)** Wing imaginal disc from *vg>GFP, PLpro, bsk-RNAi* larvae immunostained for cleaved Caspase-3 (red). **(J)** Adult wing from *vg>GFP, PLpro, bsk-RNAi* flies. Dashed lines indicate wing margins displaying the cut-wing phenotype. Scale bars: 25 μm (A, B, D, E, H, I) and 0.5 mm (C, G, J). *n* = 15 wing imaginal discs per genotype for (A, B, D–F, H, I) and *n* = 40 adult wings per genotype for (C, G, J). DAPI (blue) marks nuclei.

These features suggested activation of stress-response pathways, such as the Akt/mTOR cascade, whose chronic hyperactivation can drive metabolic overdrive and lethal oxidative stress (Nogueira et al. 2008; Los et al. 2009). Metabolic strain also intersects with JNK-dependent apoptotic signaling, creating a self-amplifying loop in which elevated ROS convert hyperactive Akt from a survival factor into a driver of epithelial cell death and tissue injury (Dhanasekaran and Reddy 2008). Notably, both Akt/mTOR and JNK pathways have been implicated in SARS-CoV-2 infection (Mukherjee and Dikic 2023; Basile et al. 2022).

In agreement, *ptc>PLpro* wing discs showed increased phosphorylation of p4E-BP (Gingras et al. 1998), a readout of Akt pathway activity (**Figure 2D**), while knockdown of Akt reduced cell death and rescued the cut-wing phenotype in PLpro-expressing tissue (**Figure 2E-G**). Conversely, constitutively active myrAkt (Stocker et al. 2002) phenocopied PLpro-induced cell death (**Supplementary Figures 4A, B**). Finally, larvae expressing *vg>PLpro* fed on food supplemented with the Akt inhibitor GSK690693 (Cheng et al. 2022) showed significant suppression of the adult cut-wing phenotype (**Supplementary Figure 4C, D**). This pharmacological rescue is in line with Guichard et al. (2023), who similarly reversed an NSP8-induced phenotype by inhibiting the relevant host-dependency factor, supporting host-pathway targeting as a general strategy for suppressing SARS-CoV-2 protein-induced perturbations (Guichard et al. 2023). Likewise, the JNK reporter TRE-dsRed was activated (**Figure 2H**) while co-expression of PLpro with *bsk-RNAi* (Lesch et al. 2010) blocked cell death in PLpro-expressing epithelium and suppressed the adult cut-wing phenotype (**Figure 2I, J**).

Taken together, these findings show that a single SARS-CoV-2 pathogenic factor, PLpro, is sufficient to uncover a conserved epithelial stress-response network comprising metabolic overdrive, oxidative stress, JNK activation, and Akt-dependent cell death in an otherwise infection-free *Drosophila* epithelium.

### PLpro-induced JAK-STAT signaling is part of a self-amplifying stress signaling network

Extensive alveolar tissue injury is a hallmark of COVID-19-associated ARDS (acute respiratory distress syndrome) (D’Agnillo et al. 2021) and is characterized by cytokine storms (Fan et al. 2026). Because injury-responsive epithelia coordinate repair through coupled JNK and JAK/STAT signaling, with JAK/STAT contributing to spatially restricted activity in the wound periphery (Quinn et al. 2025), we next asked whether PLpro engages this conserved inflammatory circuit in imaginal epithelium. In *vg>PLpro* wing epithelium, we observed a robust ∼3-fold transcriptional upregulation of the *Drosophila* IL-6 homologs *upd1, upd2, upd3* (Zandawala and Gera 2024) as well as the TNFα homolog eiger (Igaki et al. 2002) (**Figure 3A**). Consistent with this ligand induction, *ptc>PLpro* epithelial cells displayed activation of the JAK/STAT signaling reporter 10XSTAT92E-GFP in the pouch (star, **Figure 3B**). Conversely, blocking JAK/STAT signaling by co-expression of STAT-RNAi (*ptc>PLpro; STAT-RNA*i) suppressed caspase activation (**Figure 3C**) and reduced oxidative stress (DHE, **Figure 3D**), Akt signaling (4EBP, **Figure 3E**), and JNK signaling (puc-GFP, **Figure 3F**).

**Figure 3.**
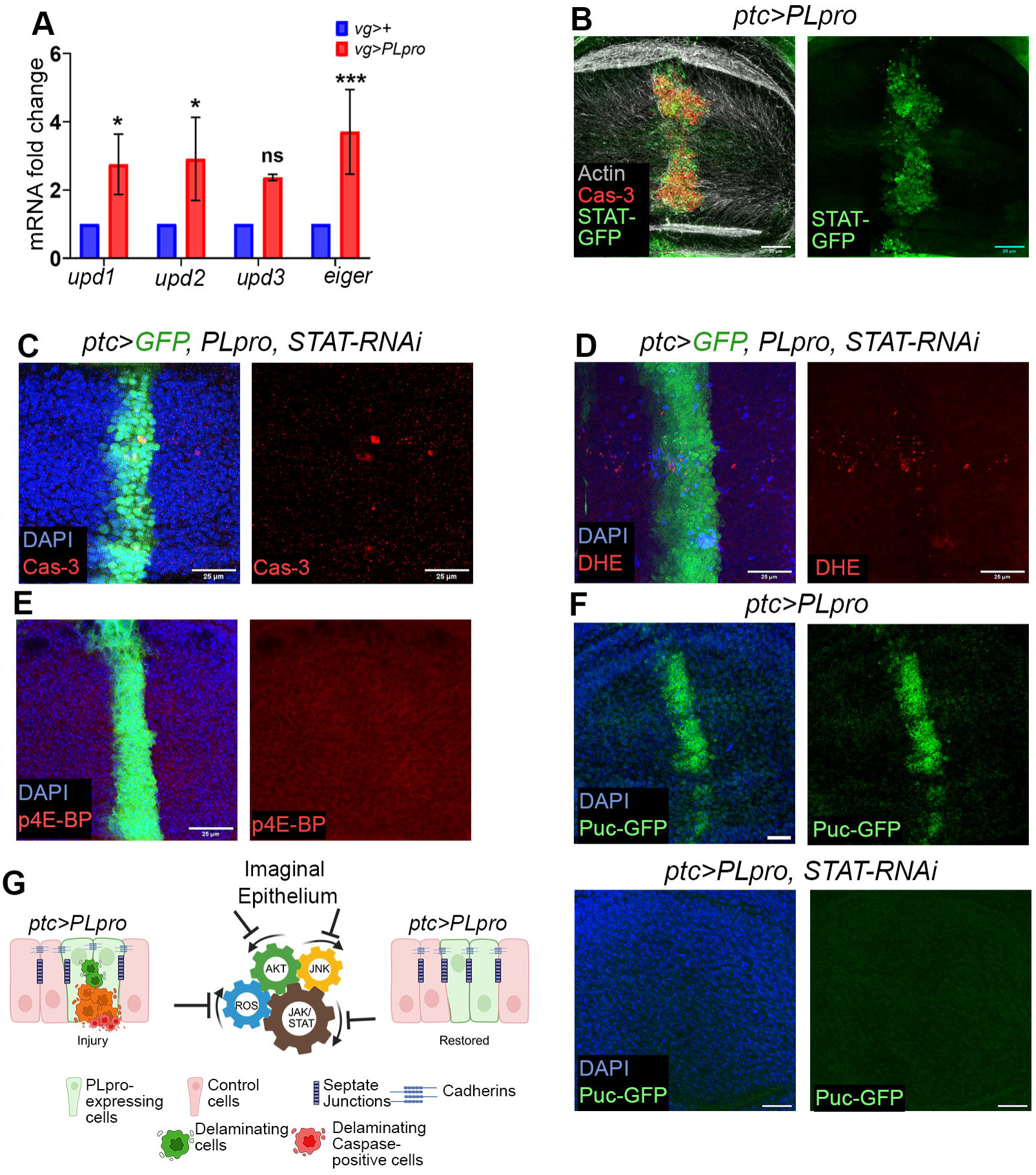
PLpro activates JAK–STAT signaling to amplify epithelial stress and tissue damage. **(A)** RT–qPCR analysis of *upd1*, *upd2*, *upd3*, and *eiger* transcript levels in wing imaginal discs from control (*vg>+*) and *vg>PLpro* larvae. Transcript abundance was normalized to the control genotype. Data are presented as mean ± SEM. *P < 0.05; ***P < 0.001; ns, not significant. **(B)** *ptc>PLpro* wing imaginal discs expressing 10XSTAT92E-GFP reporter (green), cleaved Caspase-3 staining (red), and Actin (gray). **(C–F)** *ptc>PLpro, STAT-RNAi* wing imaginal discs immunostained for cleaved Caspase-3 (red, **C**), dihydroethidium (DHE) (**D**), phosphorylated 4E-BP (p4E-BP) (**E**), and the puc-GFP reporter (JNK signaling, **F**). **(G)** Proposed model summarizing the signaling network induced by PLpro in the wing imaginal disc. PLpro activates interconnected Akt, ROS, JNK, and JAK–STAT signaling pathways that reinforce one another to promote epithelial damage. Genetic inhibition of individual pathway components disrupts this positive-feedback network and restores epithelial homeostasis. Genetic inhibition of JAK–STAT signaling suppresses multiple PLpro-induced stress responses, identifying STAT signaling as a central regulator of the pathogenic network. Scale bars: 25 μm (C–E) and 20 μm (B, F). n = 15 wing imaginal discs per genotype. DAPI (blue) marks nuclei.

A convergence of PLpro-induced cytokine production, JAK/STAT activation, JNK activation, cell death, and delamination (**Figure 1–3**) identifies a progressive inflammatory injury and tissue remodeling (Quinn et al. 2025). Rather than acting as linear cascades, these pathways form an interdependent circuit that can be reset toward homeostasis by disrupting a single signaling node, paralleling inflammatory remodeling and alveolar injury in severe COVID-19–associated ARDS (Batah and Fabro 2021). Together, these findings indicate that PLpro triggers an epithelial injury-linked inflammatory program in which JAK/STAT amplifies JNK- and ROS-dependent responses, revealing a conserved injury-to-inflammation axis that links epithelial damage to structural remodeling (**Figure 3G**).

### PLpro-induced disruption of adult intestinal epithelial barrier is JAK-STAT signaling dependent

The human gastrointestinal (GI) tract robustly expresses the ACE2 receptor, enabling ready binding to the Spike (S) proteins of SARS-CoV-2 (Lan et al. 2020), particularly in small intestinal enterocytes (Paužuolis et al. 2024). Thus, it is a major site of SARS-CoV-2 infection. COVID-19 patients often present with GI symptoms, such as diarrhea (Natarajan et al. 2022; Paužuolis et al. 2024; Xiao et al. 2020), suggesting epithelial barrier dysfunction and disrupted tight junctions (Basting et al. 2024).

Despite its relative simplicity compared to the human GI tract, the *Drosophila* adult midgut (intestine) provides a valuable model for studying gut biology (Buchon et al. 2013; Miguel-Aliaga et al. 2018). It harbors intestinal stem cells (ISCs) that differentiate into enteroblasts, enteroendocrine cells, and enterocytes, the most abundant cell type, which are characterized by large, polyploid nuclei (Zeng and Hou 2015). JAK/STAT and JNK signaling maintain ISC proliferation and homeostasis (Jiang et al. 2009; Jiang and Edgar 2011). Two well-characterized Gal4 drivers in adult midgut enterocytes (ECs), *MyoIA-Gal4* (Morgan et al. 1995) and *mex-Gal4* (Schulz et al. 1991), were used here, although *MyoIA-Gal4* shows extra-intestinal expression (Weaver et al. 2020).

*Drosophila* intestinal epithelium is sealed apicolaterally by smooth septate junctions (sSJs), forming the functional equivalent of the mammalian tight junction (Chen et al. 2020; Izumi et al. 2012). Unlike the ectodermal foregut and hindgut, which harbor pleated septate junctions (pSJs), the sSJ complex of the endodermally derived midgut comprises Snakeskin (Ssk), Mesh, and Tetraspanin 2A (Tsp2A), transmembrane proteins (Yanagihashi et al. 2012), whose mutual interdependence for sSJ localization means that perturbation of any one component reliably reflects the integrity of the entire complex (Izumi et al. 2019, 2012).

In 5-day-old *mex^ts^>PLpro* adults (expressed conditionally using a temperature-sensitive Gal80 system), intestinal ECs displayed mild cellular hypertrophy compared to controls (Ssk; **Figure 4A-C**), a sign of compensatory stress responses (Tamori and Deng 2013). By 10 days, PLpro-expressing ECs were further hypertrophied (Ssk; **Supplementary Figure 6A-C**; also see **Supplementary Figure 6D-F,** *MyoIA^ts^>PLpro*). Consistent with progressive junctional erosion, PLpro-expressing flies exhibited intestinal paracellular leakage in the “Smurf” assay — in which ingested blue dye leaks systemically upon intestinal barrier failure (Rera et al. 2012) (**Supplementary Figure 5A, B**) and a significantly shortened lifespan (**Figure 4D**), phenotypes also seen in *NSP3*-expressing flies (**Figure 4D, Supplementary 5A, B**).

**Figure 4.**
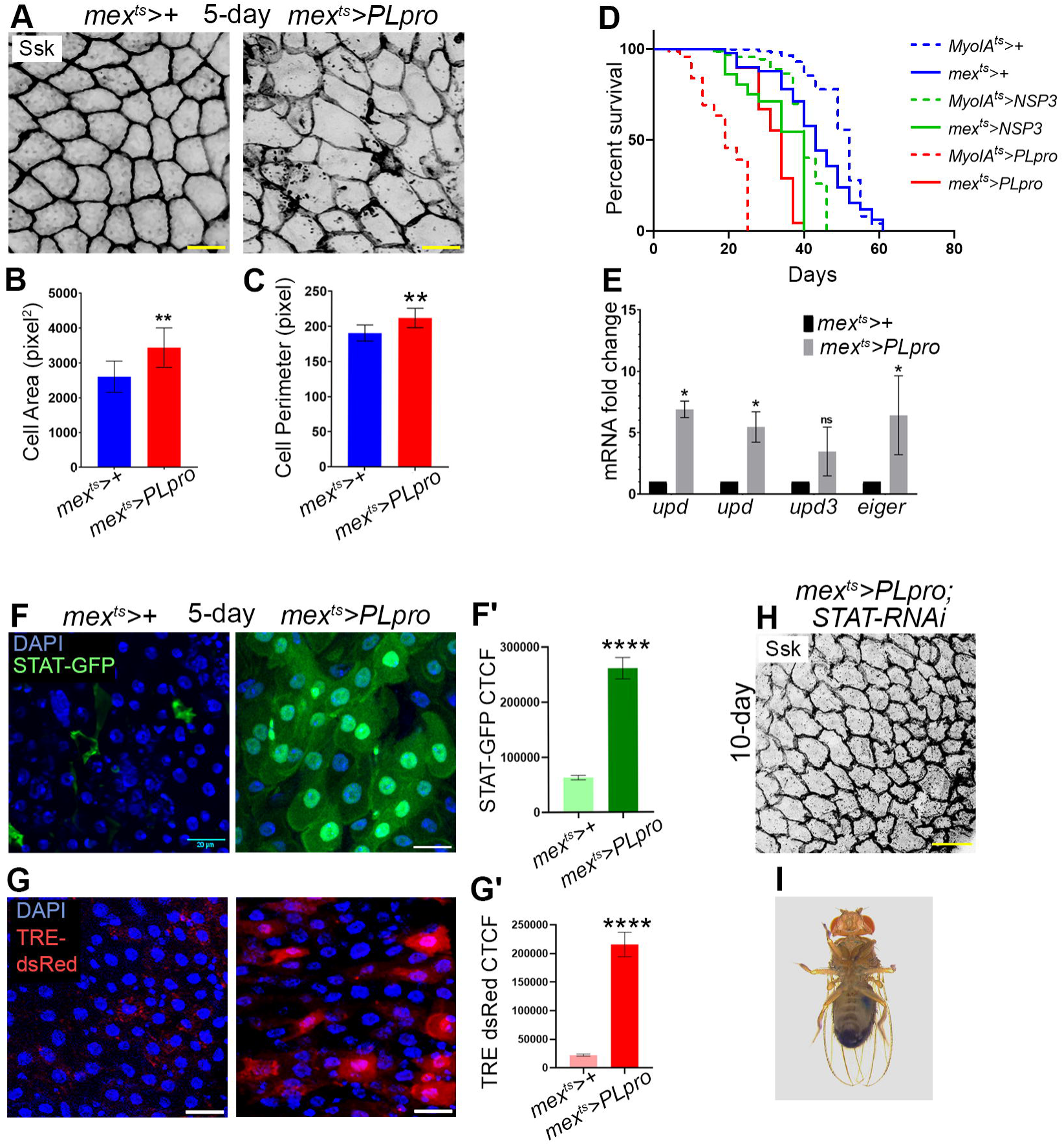
PLpro disrupts adult intestinal epithelial homeostasis by activating JAK–STAT signaling. **(A)** Posterior midgut epithelium from 5-day-old adult flies of the indicated genotypes stained for Snakeskin (Ssk; gray), a septate junction marker outlining enterocytes. **(B, C)** Quantification of enterocyte cell area (**B**) and cell perimeter (**C**) in the indicated genotypes. **(D)** Kaplan–Meier survival curves of adult flies expressing NSP3 or PLpro in enterocytes using MyoIA^ts^ or mex^ts^ drivers. Statistical significance was determined using the log-rank (Mantel–Cox) test. **(E)** RT–qPCR analysis of *upd1*, *upd2*, *upd3*, and *eiger* transcript levels in midguts of 10-day-old *mex^ts^>PLpro* adults relative to age-matched controls. **(F-F**′) Adult midguts from control (*mex^ts^>+*) and *mex^ts^>PLpro* flies expressing 10XSTAT92E-GFP reporter (green), reporter of JAK–STAT signaling. Quantification of 10XSTAT92E-GFP fluorescence intensity (**F**′). **(G-G**′) Adult midguts from control (*mex^ts^>+*) and *mex^ts^>PLpro* flies carrying the TRE-dsRed reporter (red), reporter for JNK signaling. Quantification of TRE-dsRed fluorescence intensity (**G**′). **(H)** Posterior midgut epithelium from 10-day-old *mex^ts^>PLpro, STAT-RNAi* adults stained for Snakeskin (Ssk; gray). **(I)** Smurf assay of *mex^ts^>PLpro, STAT-RNAi* adults demonstrating restoration of intestinal barrier integrity. Data are presented as mean ± SEM. ****P < 0.0001; **P < 0.01; *P < 0.05. Scale bars: 20 μm (A, F, G, H). DAPI (blue) marks nuclei in (F, G). n = 15 adults per genotype for (A, F, G, H), n = 40 adults per genotype for (I), and n = 100 adults per genotype for (D).

We next asked if, as in wing imaginal disc epithelium (**Figure 3**), perturbation of intestinal epithelial integrity is also accompanied by elevated production of inflammatory cytokines. In agreement, PLpro-expressing midguts showed transcriptional upregulation of *upd1*, *upd2*, *upd3*, and *eiger* (**Figure 4E, Supplementary Figure 6G**), mirrored by hyperactivation of the JAK/STAT reporter, *10XSTAT92E-GFP* (Bach et al. 2007) (**Figure 4F-F**′, **Supplementary Figure 6H-H**′), and the JNK reporter, *TRE-dsRed* (Chatterjee and Bohmann 2012) (**Figure 4G-G**′, **Supplementary Figure 6I-I**′). Knockdown of STAT in PLpro-expressing midgut epithelium restored epithelial architecture and reversed the *“*Smurf*”* phenotype (*mex^ts^>PLpro, STAT-RNAi,* **Figure 4H, I,** *MyoIA^ts^>PLpro; STAT-RNAi,* **Supplementary Figure 6J**).

Expression of PLpro in the *Drosophila* midgut led to a significant increase in oxidative stress and pathogenic tissue-remodeling signals, as evidenced by elevated ROS accumulation **(DHE, Supplementary Figure 7A)** and increased expression of Matrix Metalloproteases **(MMPs; Supplementary Figure 7B)** compared with control flies. These findings indicate that PLpro induces robust cellular stress and activates destructive tissue-remodeling programs in intestinal epithelial cells.

Thus, in the postmitotic enterocytes of the adult midgut, PLpro expression induces a self-amplifying host cell stress response marked by cellular hypertrophy, progressive junctional erosion, and paracellular leakage, revealing organ-intrinsic homeostatic vulnerabilities specific to the differentiated enterocyte epithelium, distinct from the injury-regeneration response of imaginal epithelia (see **Figure 1-3**).

### PLpro induces JAK-STAT signaling-dependent perturbation in tracheal cell morphology and respiration

The *Drosophila* larval respiratory system consists of two symmetrically placed dorsal trunks (DT) assembled from 10 tracheal metameres (Tr1–Tr10) (Samakovlis et al. 1996). These trunks give rise to progressively finer, anastomosing branches that terminate in tracheoles, which are functionally comparable to distal gas-exchange structures such as mammalian alveoli (**Supplementary Figure 8A**). Tracheal cell migration and branching are guided by the FGF receptor homolog Breathless (Btl) (Metzger and Krasnow 1999; Klämbt et al. 1992). The *btl-bnl* signaling, analogous to FGF/FGFR signaling in humans, regulates the finer branching pattern of the trachea (Metzger and Krasnow 1999; Sato and Kornberg 2002) and lungs (Danopoulos et al. 2019; Yang et al. 2021). Accordingly, the *btl Gal4* driver reveals GFP expression in tracheal tubes (*btl>GFP,* **Supplementary Figure 8B**). Within these tubes, chitin-lined taenidial rings maintain tracheal tube integrity (**Supplementary Figure 8A**□) (Tonning et al. 2005). Coracle—a membrane skeleton protein that marks septate junctions (Fehon et al. 1994) and DE-cadherin marks the adherens junction (Tanaka-Matakatsu et al. 1996) (**Supplementary Figure 8C**). Deep homology with mammalian respiratory systems and the genetic tractability of the *Drosophila* respiratory system have been exploited to model human respiratory diseases (Ghabrial et al. 2011; Hayashi and Kondo 2018; Roeder et al. 2012; Scholl et al. 2021).

COVID-19 lung biopsies reveal diffuse alveolar damage (DAD) characterized by AT2 cell hyperplasia, pulmonary fibrosis, infiltration of inflammatory immune cells, and ARDS hallmarks (Jyothula et al. 2022; Lamers and Haagmans 2022). Human alveolar complexity precludes full recapitulation of ARDS in *Drosophila*. Yet its tracheal epithelium offers a reductionist window into the cell-autonomous consequences of PLpro *in vivo.* We examined the effects of NSP3/PLpro expression in the *Drosophila* larval respiratory system, focusing on tracheal segment 6 (Tr6) **(Supplementary Figure 8D**) to maintain a uniform anatomical comparison across genotypes.

Late third-instar larvae actively burrow and tunnel across agar plates as they prepare for pupariation, leaving tracks reflecting locomotor and respiratory capacities (Qiang et al. 2018; Zhou et al. 2016). In this assay, *btl>NSP3* and *btl>PLpro* (**Figure 5A**) larvae both showed reduced burrowing and tunneling compared with controls, with deficits more pronounced in PLpro-expressing than NSP3-expressing larvae (**Figure 5A**). Further, compared with control tracheae, *btl>NSP3* and *btl>PLpro* tracheal cells exhibited reduced cell size (**Figure 5B** and **D**), enlarged nuclei (**Figure 5B** and **E**), and thickened Coracle-marked septate junctions (**Figure 5B** and **F**, also see **Supplementary Video 1-3**). These cells adopted a cuboidal architecture, unlike the squamous morphology of wild-type controls, indicative of epithelial thickening reminiscent of developmental gene mutations regulating tracheal tube dimensions (Ghabrial et al. 2011). A modified vertex (SI) model (Farhadifar et al. 2007) further supported that PLpro-induced tracheal geometry defects are cell-intrinsic (**Supplementary Figure 9A, B**). Chitin, essential for tracheal lumen diameter organization and tube morphogenesis (**Figure 5C**) (Luschnig et al. 2006; Tonning et al. 2005), displayed loss of regular concentric luminal lining in both NSP3- and PLpro-expressing tracheae (**Figure 5C**). Beyond these structural consequences, NSP3- and PLpro-expressing tracheal epithelia—like their imaginal (Figures 1-2) and intestinal (**Figure 4**) counterparts—displayed hypoxia (LDH-lacZ; **Supplementary Figure 10A)** (Lavista-Llanos et al. 2002), and heightened JNK activity (TRE-dsRed; **Supplementary Figure 10B**; (Chatterjee and Bohmann 2012)).

**Figure 5.**
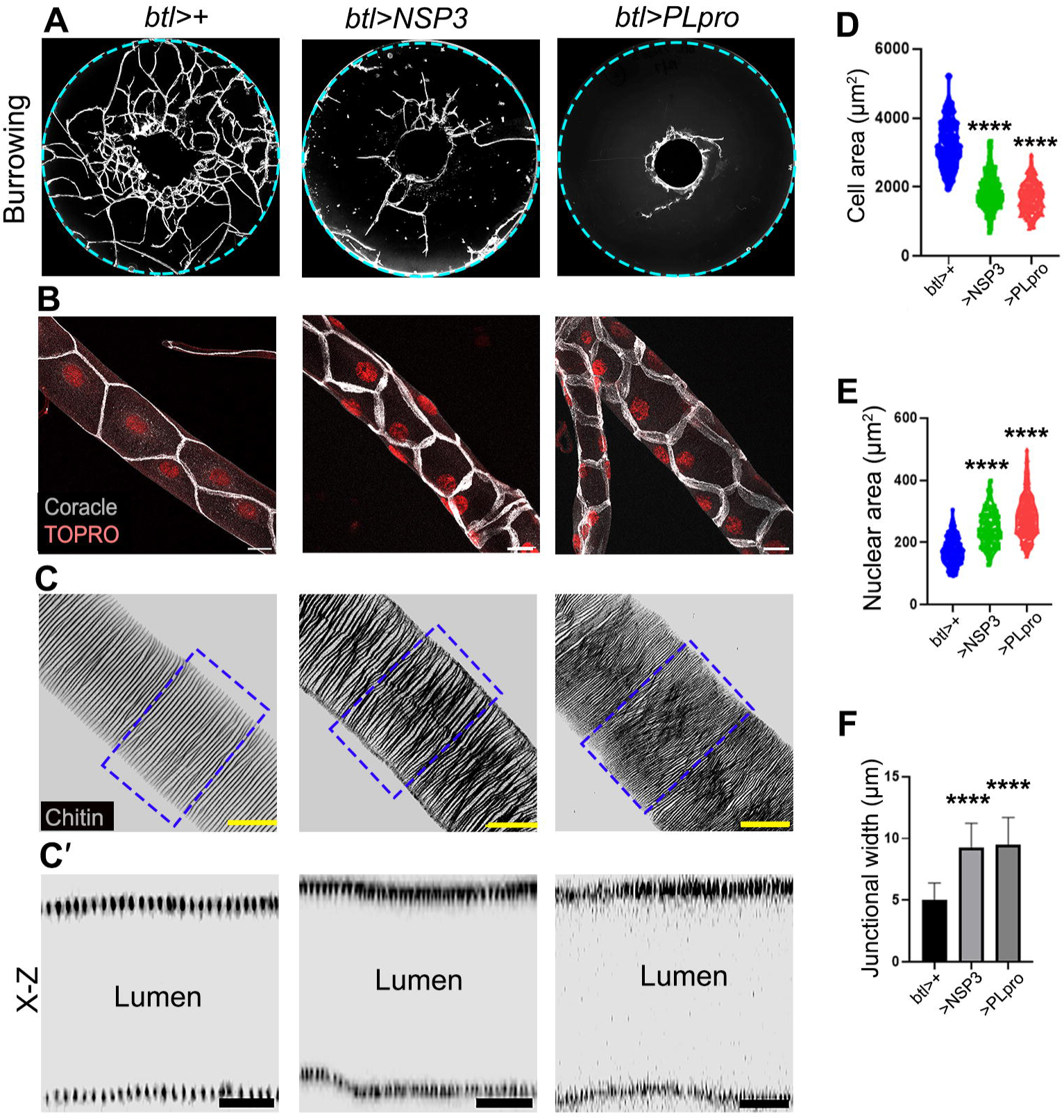
PLpro induces a squamous-to-cuboidal transition in dorsal tracheal trunk cells and promotes larval hypoxia. **(A)** Agar plates from burrowing assays of third-instar *btl>+* (control), *btl>NSP3*, and *btl>PLpro* larvae. The light blue dashed line marks the perimeter of the Petri dish. **(B)** Dorsal tracheal trunks (DTs) from the indicated genotypes immunostained for Coracle (gray). **(C–C**′) DTs from the indicated genotypes stained for chitin (gray). (C′) X–Z optical sections of the boxed regions shown in (C). **(D-F)** Quantification of tracheal cell area (**D**), nuclear area (**E**), and cell junction width (**F**) of the indicated genotypes. Nuclei are counterstained with TO-PRO (red) in (B). Data are presented as mean ± SEM. Statistical significance was determined using one-way ANOVA. ****P < 0.0001. Scale bars: 20 μm (B, C) and 10 μm (C′). n = 10 larvae per genotype for (A), n = 15 larvae per genotype for (B–C′), and n > 200 cells per genotype for (D–F).

Finally, like their larval wing (**Figure 3**) and adult intestinal (**Figure 4**) epithelial counterparts, *PLpro*-expressing tracheal tubes also displayed upregulated JAK-STAT signaling (10XSTAT92E-GFP; **Figure 6A**) (Bach et al. 2007). Knockdown of JAK-STAT in NSP3-expressing trachea improved respiratory health *(btl>NSP3, STAT*-*RNAi;* **Figure 6B**), with reversal of cuboidal to flattened morphology, restoration of narrow Coracle bands at septate junctions (**Figure 6C, E-G**, *also see* **Supplementary Video 4**), and rescued chitin luminal distribution (**Figure 6D, D□**).

**Figure 6.**
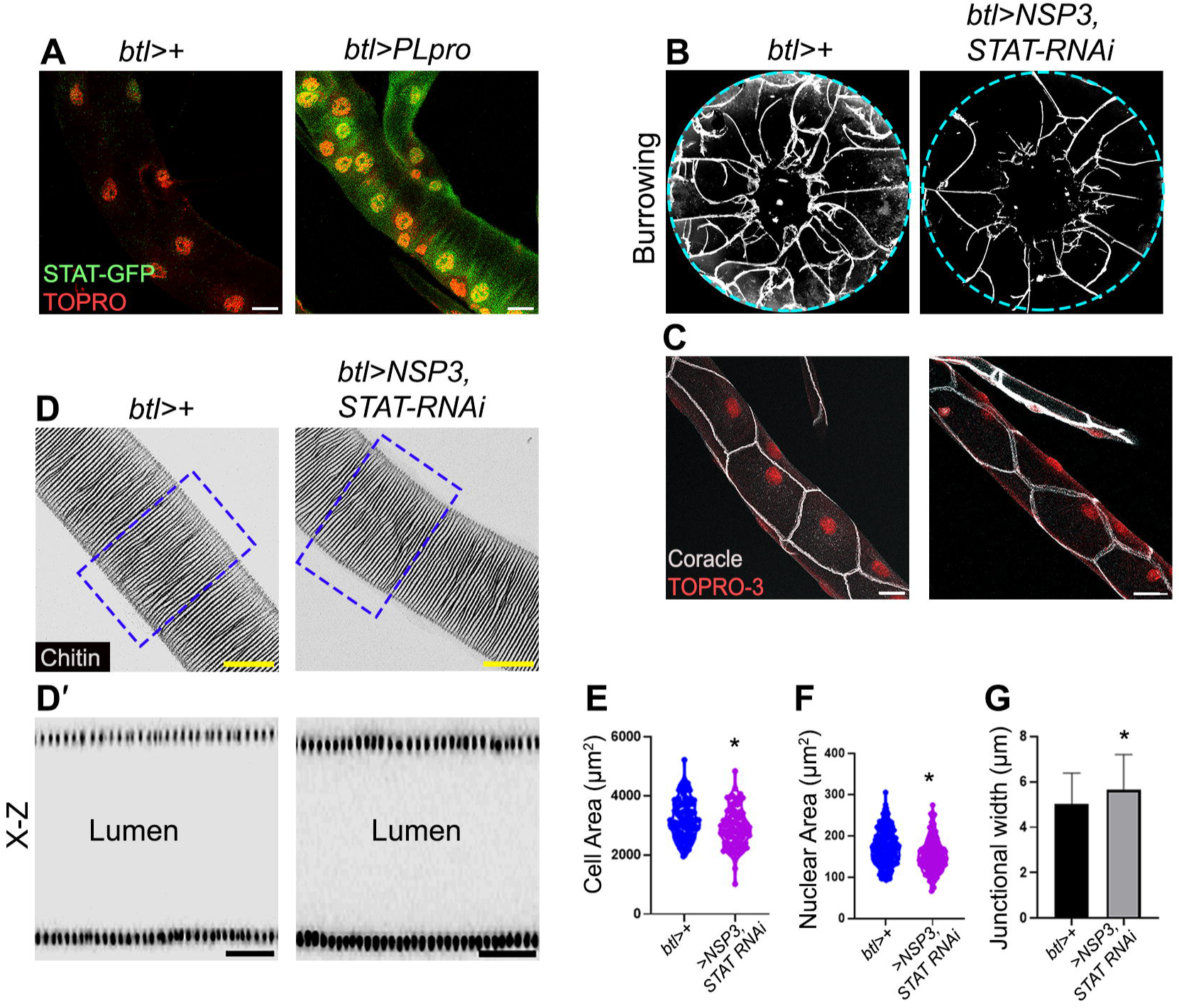
PLpro-induced tracheal remodeling and hypoxia are STAT-dependent. **(A)** Larval dorsal tracheal trunks (DTs) from *btl>+* (control) and *btl>PLpro* larvae expressing the STAT-GFP reporter (green). **(B)** Agar plates from burrowing assays of *btl>+* (control) and *btl>NSP3, STAT-RNAi* larvae. The light blue dashed line marks the perimeter of the Petri dish. **(C)** Larval DTs from the indicated genotypes stained for Coracle (gray). **(D–D**′) Larval DTs from the indicated genotypes stained for chitin (gray). (D′) X–Z optical sections of the boxed regions shown in (D). **(E–G)** Quantification of tracheal cell area (**E**), nuclear area (**F**), and Coracle-positive cell junction width (**G**) in the indicated genotypes. Data are presented as mean ± SEM. *P < 0.05. Scale bars: 20 μm (A, C, D) and 10 μm (D′). TO-PRO-3 (red) marks nuclei in (A, C). n = 10 larvae per genotype for (B), n = 15 larvae per genotype for (A, C–D), and n > 200 cells per genotype for (E–G).

In essence, PLpro-induced squamous-to-cuboidal transition, thickened septate junctions, and disrupted chitin architecture in dorsal tracheal trunks resemble fibrotic airway remodeling in COVID-19 and long COVID (Jyothula et al. 2022), indicating that PLpro drives respiratory epithelial remodeling in a cell-autonomous manner (Huizen et al. 2024).

### Hypersensitivity of tracheoblasts and adult air-sac precursor cells to NSP3/PLpro-driven JAK-STAT signaling

During metamorphosis, multiple progenitor sources remodel the larval tracheal system to generate the adult respiratory network. The wing-disc-associated air sac precursor (ASP) cells (**Figure 7A, B**) give rise to the adult dorsal air sacs that oxygenate the adult flight muscles (Guha and Kornberg 2005; Guha et al. 2008; Sato and Kornberg 2002). Separately, Tr2 tracheoblasts (**Figure 7A, B**) are proliferation-competent adult progenitors that contribute to adult tracheal remodeling and organogenesis (Guha et al. 2008; Li et al. 2022; Pitsouli and Perrimon 2010). These progenitor pools are functionally linked through the transverse connective, which connects the Tr2 lineage to ASP morphogenesis. The formation of the ASP depends on the proliferation, migration, and viability of Tr2-derived tracheal cells (Guha and Kornberg 2005). Importantly, these progenitor pools are controlled by Bnl/FGF signaling, whose genetic perturbation abrogates ASP growth and Tr2 tracheoblast-driven remodeling (Guha and Kornberg 2005; Guha et al. 2008; Li et al. 2022; Pitsouli and Perrimon 2010; Sato and Kornberg 2002). This remodeling is regionally specified: distinct tracheoblast pools contribute to different thoracic and abdominal tracheal derivatives rather than following a single, linear precursor-to-structure mapping (Ghabrial et al. 2011; Metzger and Krasnow 1999; Pitsouli and Perrimon 2010).

**Figure 7.**
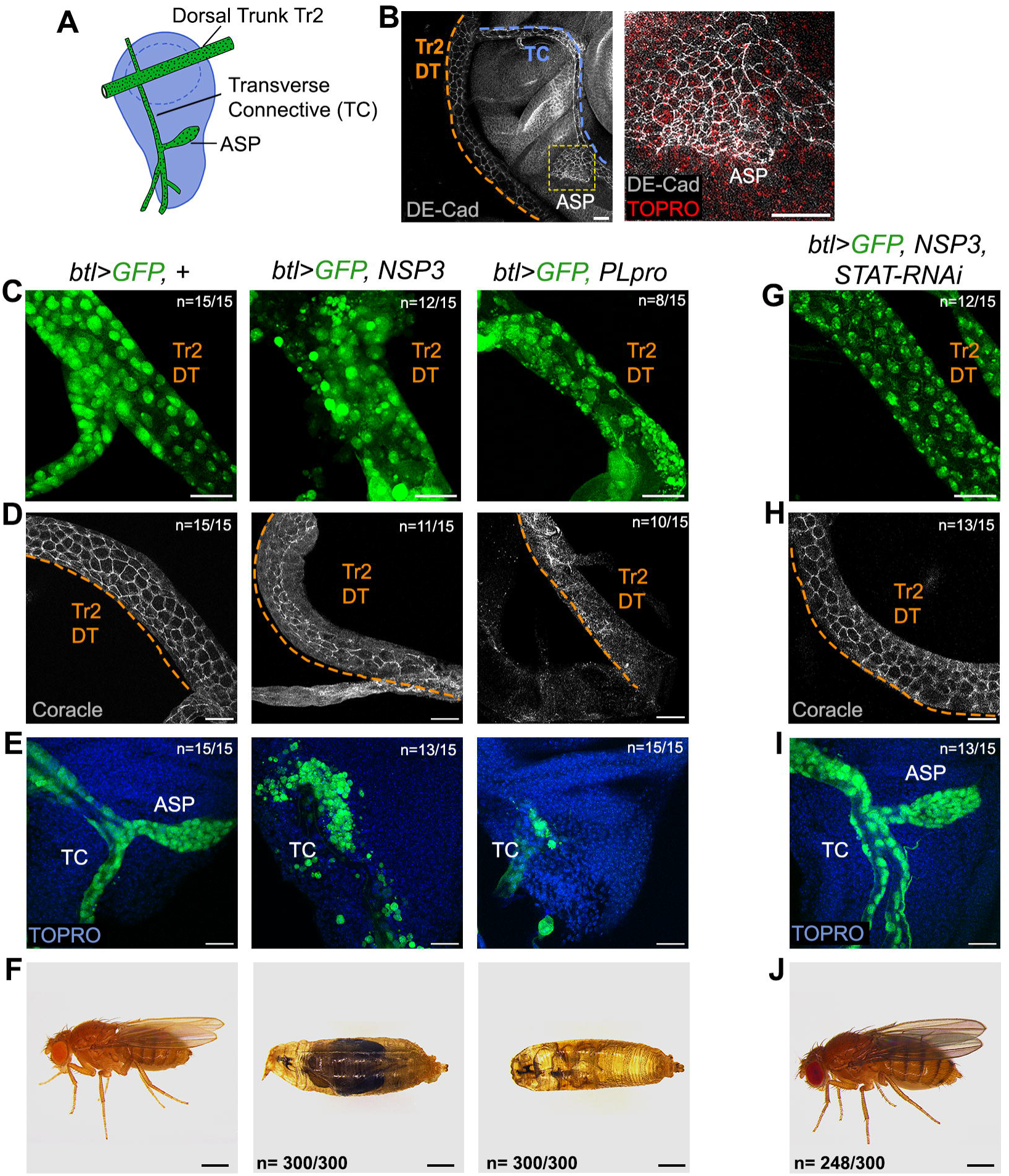
PLpro disrupts tracheoblast maintenance, air sac primordium development, and adult viability through STAT-dependent signaling. **(A)** Schematic representation of the second dorsal trunk metamere (Tr2) of the late third-instar larval trachea. The Tr2 region contains tracheoblasts that give rise to the air sac primordium (ASP) and is connected to the wing imaginal disc through the transverse connective (TC). **(B)** Late third-instar larval preparation showing the Tr2 region, transverse connective (TC), and ASP associated with the wing imaginal disc. DE-cadherin (gray) marks tracheal epithelial cells. The right panel shows a higher magnification of the boxed region. **(C–E)** Tr2 region from *btl>GFP, +* (control), *btl>GFP, NSP3*, and *btl>GFP, PLpro* larvae showing GFP-positive tracheoblasts (**C**), Coracle staining (gray) (**D**), and TO-PRO-3-labeled nuclei (**E**). **(F)** Adult flies and pharate adults of the indicated genotypes. **(G–I)** Tr2 region from *btl>GFP, NSP3, STAT-RNAi* larvae showing GFP-positive tracheoblasts (**G**), Coracle staining (gray) (**H**), and TO-PRO-3-labeled nuclei (**I**). **(J)** Adult flies from *btl>GFP, NSP3, STAT-RNAi*. Scale bars: 20 μm (B–E, G–I) and 0.5 mm (F, J). TO-PRO-3 marks nuclei in (B, E, I). n = 20 larvae per genotype for (B–E, G–I) and n = 300 adults per genotype for (F, J).

Consistent with broad disruption of tracheal architecture and respiratory function (**Figures 5** and **6**), *btl>NSP3, GFP* and *btl>PLpro*, *GFP* larvae showed marked depletion of Tr2 tracheoblasts (**Figure 7C**) and epithelial disruption (Coracle, grey) relative to *btl>GFP* controls (**Figure 7D**). PLpro expression disrupted Tr2 remodeling, producing conspicuous gaps in the second thoracic tracheal lineage and consistent with failure of the progenitor-derived tracheal program. Correspondingly, wing imaginal discs displayed near-complete loss of both the transverse connective and ASP cells in both *btl>NSP3, GFP* and *btl>PLpro, GFP* larvae (**Figures 7E**). Because ASP morphogenesis depends on both proliferation and migration (Cruz et al. 2015; Guha and Kornberg 2005; Guha et al. 2008), loss of viable transverse connective cells likely accounts for the failure of ASP outgrowth. Both *btl, GFP>NSP3* and *btl>PLpro, GFP* larvae failed to complete metamorphosis and died as pseudo-pupae (**Figure 7F**). Finally, knockdown of JAK-STAT signaling rescued epithelial integrity in Tr2 (**Figure 7G, H**), restored repopulation of the transverse connective with ASP formation (**Figure 7I**), and rescued pupal lethality, thereby restoring adult eclosion in NSP3-expressing larvae (**Figure 7J**).

Together (**Figure 5-7**), these findings show that differentiated tracheal tubes are compromised by PLpro-induced JAK-STAT signaling, whereas the progenitor pools are severely depleted. Human lung epithelium likewise contains epithelial lineages with distinct renewal capacities (Rawlins 2008) and differential susceptibility to SARS-CoV-2 infection and severe COVID-19 (Mulay et al. 2021; Wang et al. 2022; Ting et al. 2022). Notably, genetic inhibition of JAK-STAT signaling rescued PLpro-induced respiratory distress in flies. This targeted rescue parallels the clinical efficacy of specific JAK1/2 inhibitors, such as baricitinib, which have been successfully deployed in severe COVID-19 to attenuate cytokine-driven hyperinflammation and improve patient survival (Gu et al. 2025; Dupuis et al. 2022) and identifying JAK-STAT signaling as a tractable therapeutic node.

### PLpro induces junctional condensation and incipient fibrotic remodeling in mammalian MDCK epithelia

We next asked whether the epithelial defects induced by PLpro in *Drosophila* are conserved in mammalian epithelia. MDCK (Taub et al. 1979) cells form a well-defined epithelium *in vitro*, marked by sub-apical tight junctions (TJ) (Fu et al. 2022; Otani and Furuse 2020), analogous to lateral *Drosophila* SJ (Nelson et al. 2010; Willott et al. 1993) and apico-lateral AJ (Itoh et al. 1997), as in insect epithelium (Bilder 2004) (**Figure 8A**). To test PLpro activity in this context, we generated MDCK cells stably expressing mammalian codon-optimized, Myc-tagged PLpro under tetracycline control (**Supplementary Figure 11A**). We assessed expression using the PLpro antibody and observed robust induction upon doxycycline treatment (**Supplementary Figure 11B**).

**Figure 8.**
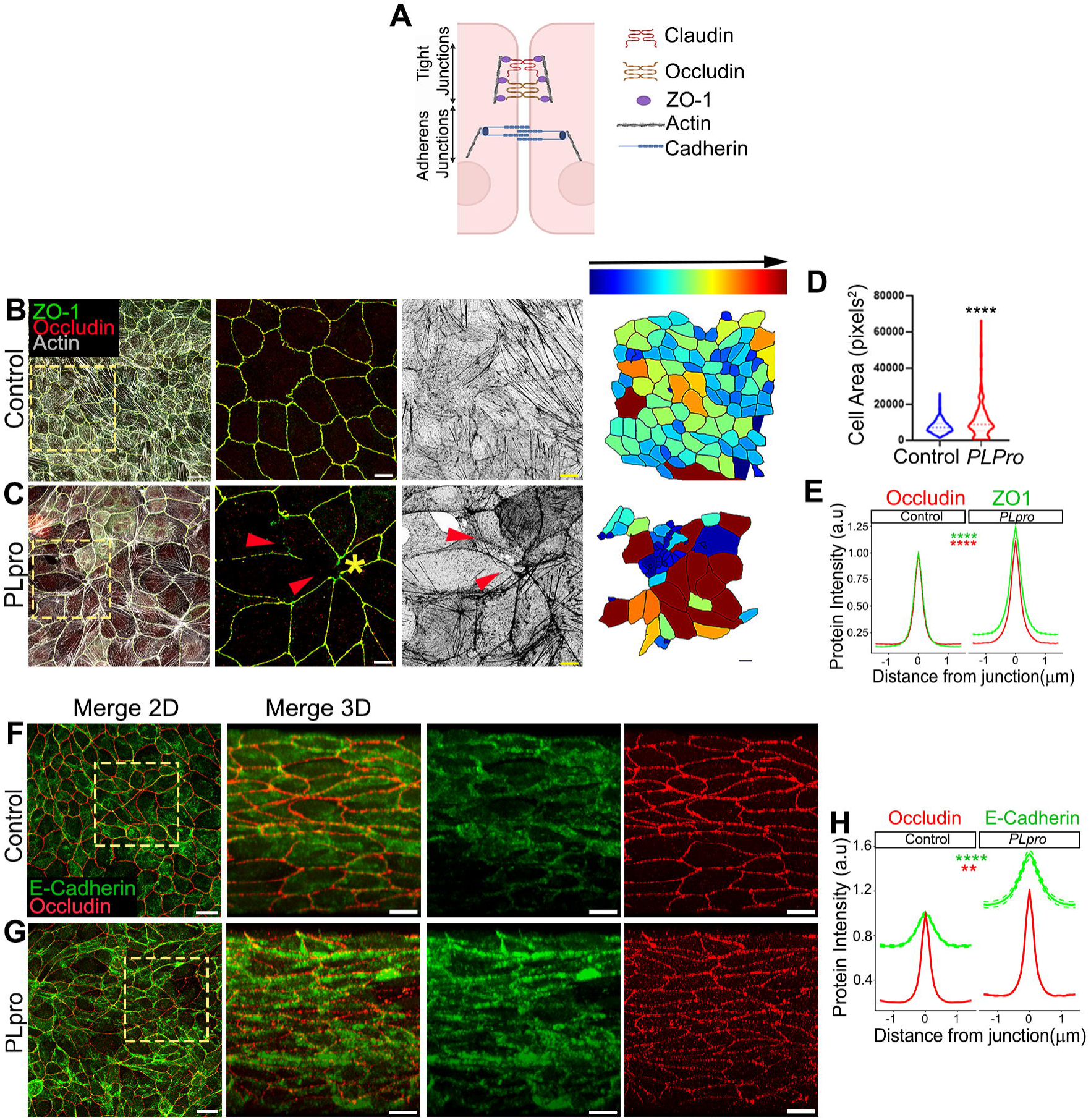
PLpro induces cytoarchitectural and tight-junctional damage and cellular stress in MDCK cells. **(A)** Schematic of mammalian epithelial junctions and cytoskeletal proteins examined in this study, including the tight junction components Zonula Occludens-1 (ZO-1) and Occludin, the adherens junction marker E-cadherin, and filamentous actin (F-actin). **(B–D)** Immunofluorescence images of ZO-1 (green), Occludin (red), and F-actin (Phalloidin, gray) in control (**B**) and PLpro-expressing (**C**) MDCK cells. Boxed regions are shown at higher magnification in the adjacent panels. Segmented images pseudocolored with the jet colormap were used for quantification of cell area (**D**). **(E)** Quantification of ZO-1 and Occludin fluorescence intensities across cell junctions. **(F–G)** Two-dimensional (X–Y) and three-dimensional projections of E-cadherin (green) and Occludin (red) in control (**F**) and PLpro-expressing (**G**) MDCK cells. Individual 3D projections of E-cadherin and Occludin from the boxed regions are shown in the right panels. **(H)** Quantification of E-cadherin and Occludin fluorescence intensities across cell junctions. Data are presented as mean ± SEM. ****P < 0.0001; **P < 0.01. Scale bars: 20 μm (B–C, F–G, first panels) and 10 μm (B–C, F–G, remaining panels). n > 300 junctions per condition for (D, E, H). n = 5 independent experiments.

After 72 h of doxycycline induction, *PLpro*-expressing cells showed disrupted ZO-1 and Occludin organization at tight junctions compared with controls (**Figure 8C** versus **Figure 8B**). Perijunctional actin remained evident despite junctional disorganization, and the epithelium contained rosette-like cell clusters with altered cell shape (**Figure 8C**), suggestive of apical extrusion (Kuipers et al. 2014; Teo et al. 2020). Fluorescence image segmentation analysis (SI) showed increased mean cell area and greater cell-size variability in PLpro-expressing epithelia relative to controls (**Figure 8B-D**). Junctional ZO-1 and Occludin intensity was also increased (**Figure 8E**), indicating condensation and redistribution rather than simple loss of tight junction components. E-cadherin is a core determinant of adherens junctions (AJs) in polarized epithelia and is essential for cell-to-cell adhesion (Coopman and Djiane 2016). AJ formation precedes TJ formation, and is required for its maturation (Shigetomi et al. 2018) while actomyosin tension generated at AJs stabilizes TJ barriers (Coopman and Djiane 2016; Itoh et al. 1997; Shigetomi et al. 2018). 2- and 3-dimensional projections (**Figure 8F, G**) of AJ (E-cadherin, green; (Coopman and Djiane 2016)) and TJ (Occludin, red,(Cummins 2012)) revealed irregular junctional continuity and cytoplasmic mislocalization of E-cadherin (**Figure 8G**). Quantification of junctional E-Cadherin and Occludin intensity further confirmed these perturbations (**Figure 8H**). Together, these data indicate remodeling of both adherens and tight junctions in response to PLpro.

*PLpro*-expressing MDCK cells also exhibited elevated ROS, increased DNA damage, and more cell death (**Supplementary Figure 11C-E**□). Finally, PLpro induced transcriptional upregulation of IL-6 and STAT5 (**Supplementary Figure 11F**), extending the inflammatory signaling phenotype observed in *Drosophila* to a mammalian epithelial system.

## DISCUSSION

At its core, COVID-19 manifests as a systemic epithelial disorder in which major clinical complications, such as acute respiratory distress syndrome (ARDS) and gastrointestinal dysfunction, arise from compromised epithelial integrity (Duan et al. 2024; Davis et al. 2023; Deinhardt-Emmer et al. 2021). Large-scale efforts to assign pathogenic roles to individual SARS-CoV-2 proteins have yielded important interaction maps and cell-based phenotypes (Hayn et al. 2021; Zhou et al. 2023; Gordon, Hiatt, et al. 2020; Miao et al. 2021). However, these studies have not directly resolved how individual viral factors drive loss of epithelial homeostasis in intact tissues (Becker et al. 2024; Ackermann et al. 2020; Basting et al. 2024). Against this background, the present study uses an unbiased in vivo screen to reveal that NSP3-PLpro, as a SARS-CoV-2 factor, is sufficient to induce epithelial architectural disruption, cell-autonomous inflammatory signaling, and cell death across multiple epithelial organs in *Drosophila*, even in the absence of viral infection. The early increase in glucose uptake, ROS accumulation, oxidative DNA damage, and hyperactivation of Akt, JNK, and JAK/STAT, together with caspase activation and basal delamination, supports the view that JAK/STAT is part of the injury-amplification loop. Its suppression therefore reverses much of the associated stress signaling, tissue damage, and cell death in PLpro-expressing epithelia. Further, its activation in PLpro-expressing mammalian MDCK epithelia with junctional remodeling suggests that this logic of epithelial injury is conserved across phylogeny (Ezeonwumelu et al. 2021; Sarapultsev et al. 2023). Thus, while PLpro has previously been defined by its roles in SARS-CoV-2 polyprotein processing and antagonism of host innate immunity through its DUB/deISGylase activities (Shin et al. 2020; Cao et al. 2023; Ullrich and Nitsche 2022), our findings reveal an additional pathogenic activity in structurally intact epithelia in an infection free context.

PLpro expression in the *Drosophila* larval respiratory system reveals compartment-specific vulnerability. In differentiated tracheal tubes beyond the Tr2 lineage, PLpro drives squamous-to-cuboidal remodeling, septate-junction thickening, and disruption of the ordered luminal chitin architecture, producing a phenotype consistent with maladaptive epithelial remodeling rather than nonspecific degeneration (Wagner et al. 2021). The reversal of these perturbations by JAK-STAT knockdown suggests that the remodeling is inflammation-driven and spatially restricted. In this respect, PLpro-induced tracheal defects resemble a fibrosis-like epithelial response, in which epithelial architecture is remodeled in association with persistent injury signaling (Kalluri and Weinberg 2009; Ehrhardt et al. 2022; Wagner et al. 2021). This compartmentalization is also consistent with the broader principle that airway epithelial JAK-STAT activity is required for homeostasis and stress resilience in *Drosophila*, whereas excessive signaling drives cell-autonomous remodeling and loss of epithelial homeostasis (Niu et al. 2026).

Tracheoblasts of the Tr2 metamere and the transverse connective are a developmentally plastic progenitor population that depends on Branchless-FGF signaling delivered through actin-based cytoneme contacts for growth and morphogenesis (Sato and Kornberg 2002; Roy et al. 2011; Du et al. 2022). In this setting, PLpro likely compromises the progenitor niche by disrupting epithelial architecture, cortical actin organization, junctional stability, and cell survival, thereby weakening the cytoneme-dependent signaling environment required for air sac primordium (ASP) maintenance and fate progression (Guha and Kornberg 2005; Du et al. 2018). That PLpro-induced tracheoblast depletion in the Tr2 tracheal tube and transverse connective, or the loss of ASP, is reversed by knockdown of JAK-STAT signaling, further suggests its perturbation of cellular signaling(s) that maintains the tracheal progenitor cell fate. Tracheoblast identity is maintained by a Notch-dependent transcriptional program that includes regulators such as cut and blistered (Li et al. 2024), indicating that the progenitor compartment is not simply less differentiated tissue, but a distinct developmental state requiring active maintenance (Guha et al. 2008; Guha and Kornberg 2005). Against this background, PLpro-induced JAK-STAT activity may destabilize a Notch-supported tracheoblast program, thereby compromising progenitor maintenance. Our observation in the larval tracheal system therefore is reminiscent of other *Drosophila* progenitor systems in which Notch and JAK/STAT exert opposing influences on cell fate and proliferation (Bausek 2013; Liu et al. 2010). Finally, PLpro-expressing tracheal cells are also likely to destabilize the nuanced JAK-STAT signaling required for tracheal cell homeostasis (Niu et al. 2026).

Taken together, PLpro appears to impact the two respiratory compartments through a common inflammatory axis: in differentiated tubes, it promotes epithelial thickening and architectural remodeling, whereas in the progenitor-like Tr2/air sac primordium compartment, it disrupts the signaling and structural conditions required for progenitor survival and maintenance.

In the healthy lung, resident epithelial stem and progenitor cells—such as airway basal cells and alveolar type 2 (AT2) cells—maintain tissue homeostasis and drive regeneration after injury (Bi et al. 2025; Rawlins 2008; Tripathy et al. 2026). In severe COVID-19 lungs, transient progenitors fail to terminally differentiate, driving maladaptive fibroproliferative remodeling (Ting et al. 2022; Jyothula et al. 2022; Lamers and Haagmans 2022). Crucially, our *Drosophila* model demonstrates that this pathogenic mechanism is evolutionarily conserved, where triggers like PLpro hijack the host JAK-STAT pathway to lock the respiratory epithelium into a persistent, fibrotic state. Consequently, we propose that the clinical efficacy of JAK-STAT inhibitors in improving severe ARDS outcomes (Dupuis et al. 2022; Gu et al. 2025) is mechanistically akin to the RNAi-mediated restoration of tracheoblasts and air sac primordium (ASP) populations observed in our fly model.

These findings also provide a rationale for host-directed intervention. Although current efforts have focused on direct PLpro inhibition (Bader et al. 2025; Tan et al. 2024; Jadhav et al. 2025; Lu et al. 2024; Garnsey et al. 2024), interruption of host nodes such as Akt or JAK/STAT may likewise be sufficient to collapse the broader injury circuit, consistent with host-directed mitigation of severe COVID-19 (Stebbing et al. 2021; Gu et al. 2025; Dupuis et al. 2022). Finally, by revealing that Plpro triggers a self-amplifying epithelial stress circuit, our findings also point to a shared logic with non-viral diseases such as cancer (Galbraith et al. 2023; Taniguchi et al. 2020; Piao et al. 2022) and metabolic disorders (Dong et al. 2024; Mo et al. 2025), where sustained stress signaling, metabolic rewiring, and tissue remodeling drive progressive pathology. In this context, the *Drosophila* model, given its capacity to uncover viral-host interactomes (Guichard et al. 2023), offers a rapid platform for host-targeted drug discovery.

## Caveats and future directions

Our data establish PLpro as a pathogenic factor but do not yet resolve which of its enzymatic activities is mechanistically responsible for the epithelial injury phenotype, namely, its deubiquitinating, cysteine protease, or de-ISGylation activity. Because *Drosophila* **l**acks the canonical RIG-I/MDA5–MAVS–IRF3/7 pathway that drives type I interferon production (Cao et al. 2023; Lemaitre and Hoffmann 2007), the contribution of PLpro’s de-ISGylating activity to the epithelial defects described here is likely to be limited, or at least not directly testable in this system. Notably, selective ablation of PLpro’s DUB activity did not alter viral replication or innate immune responses *in vivo* (Huizen et al. 2024), arguing against an obligate requirement for this activity in all major PLpro-dependent pathogenic outputs. Taken together, these considerations narrow the mechanistic possibilities but do not distinguish whether epithelial injury is driven predominantly by one catalytic function or by cooperative contributions from more than one. Future work using catalytically selective PLpro mutants will therefore be required to identify the activity that is necessary and sufficient for epithelial injury, while preserving the central conclusion of this study that PLpro is itself sufficient to trigger it.

## Materials and Methods

Fly lines were cultured on standard yeast-cornmeal agar at 25°C. All the genetic stocks and reagents used in this study were obtained from public repositories or as gifts from other researchers (SI). q-PCR to assess the regulatory status of candidate cytokines in three *Drosophila* epithelia and MDCK cells was performed using standard protocols; primers used are listed in the SI. Immunohistochemical staining was performed by dissecting tissues in PBS, fixing them in 4% PFA, and staining with the desired antibody. A detailed description is given in SI.

## Data, Materials, and Software Availability

All study data are included in the article and/or supporting information (SI). The codes used in the study for junctional segmentation in gut and MDCK cells, and for tracheal modeling, are available at https://github.com/suhailrbme/tracheaAnalysisModel.git.

## Supporting information

Suppmentary Information (SI)

## Acknowledgment

We acknowledge IIT Kanpur for its support for research in the PS laboratory; DBT, MHRD, UGC, and CSIR for financial support to QTA, SB, IMA, JT, SSP, and AB; and the DST-KIRAN WOS-A fellowship, which additionally supported AB. We thank Nitin Mohan for providing access to his cell culture facility. Authors thank Arjun Guha for many rounds of critical discussion on tracheal phenotypes.

## Author contributions

QTA, SB, and PS conceived and designed the study. QTA characterized the wing imaginal epithelium, while SB characterized the tracheal and MDCK epithelia. IA, JT, and QTA characterized the intestinal epithelium. AB and SSP contributed to data analysis and validation. MDSR and SB performed fluorescence intensity quantification and tracheal cell modeling. QTA, SB, and PS wrote the manuscript, and all authors contributed to editing and validation of the final version.

