## Supplementary material for "SARS-CoV-2 PLpro Hijacks a Conserved Stress Signaling Network to Drive Organ-Specific Loss of Epithelial Homeostasis": Suppmentary Information (SI)

**Keywords**

This PDF file includes:

A. Supplementary Text
B. Supplementary Table (S1-S5)
C. Supplementary References
D. Supplementary Figure legends (Figure 1-11)
E. Supplementary Video legends (1-4)

**SUPPLEMENTARY INFORMATION, SI**

**EXPERIMENTAL METHODS**

**Genetic screens of SARS-CoV-2 nonstructural genes**

Two sources of transgenic *Drosophila* lines were used in this study. First, the entire *Drosophila* COVID-19 Resource (DCR) (Table S1, (Guichard et al. 2023)), obtained from BDSC, each representing an individual SARS-CoV-2 protein-coding gene. We selectively screened the nonstructural and accessory protein-coding transgenes from this collection. In this screen, NSP3 expression in larval wing epithelium induced a prominent cut-wing phenotype (**Fig. S1**), which prompted our further subcloning of its PLpro domain (source of NSP3, Addgene 141257 pDONR207 SARS-CoV-2 NSP3) via PCR amplification, restriction digestion, sequence verification and subcloning in *Drosophila* pUASTattB vector (Bischof et al. 2007) and, finally, transgenesis via φC31-based targeted integration at ZH-attP-86Fb (BDSC stock # 24749). Transgenesis was carried out at the C-CAMP Fly (*Drosophila*) Facility, Bengaluru. *PLpro* expression was then examined in multiple epithelial organs using specific Gal4 drivers using Gal4/UAS binary system (Brand and Perrimon 1993).

*Drosophila* cultures were reared on standard cornmeal agar food at 25±0.5°C. The Gal4 lines used for this work are boundary enhancer-driven *vg-Gal4* (Simmonds et al. 1998), *ptc-Gal4* (BDSC), and *en-Gal4* (Simmonds et al. 1995) for larval wing imaginal disc-specific expression. For adult mid-gut specific expressions, *MyoIA Gal4; tub Gal80^ts^* (Jiang et al. 2009; Morgan et al. 1994) and *mex-Gal4; tub Gal80^ts^* (Marchetti et al. 2022; Schulz et al. 1991) drivers were used, while *btl-Gal4* (Shiga et al. 1996) was used for tracheal expression. Candidate transgenes were expressed under the *MyoIA-Gal4* or *mex-Gal4* driver in a *tub-Gal80^ts^* genetic background (shortened as *MyoIA-Gal4^ts^* and *mex-Gal4^ts^*). Larval cultures were thus reared at 18°C until adult emergence, then shifted to 29°C for adult mid-gut-specific expression of the candidate transgene.

**Adult wing phenotypic study and quantifications**

Flies were separated by sex and preserved in 70% (v/v) ethanol for at least 24 hrs. Flies with damaged or crumpled wings were excluded before mounting. Wings were mounted using the established protocol (Alba et al. 2021) and imaged under brightfield illumination with a Leica M205 FA stereomicroscope. Wing area was quantified in FIJI using the freehand drawing tool to trace the boundaries of wild-type and NSP-expressing transgenic adult wings. For transgenic wings showing a cut-wing margin, a hypothetical line was drawn to reconstruct the presumptive intact wing. The area of wing margin loss was calculated by subtracting the measured area of transgenic wings from the corresponding reconstructed presumptive wing area. Statistical analysis and graphical representation were performed using GraphPad Prism 8.0; a threshold of *p*<0.05 was used to define statistical significance.

**Immunofluorescence and image analysis**

Third-instar larval tissues were dissected in PBS and fixed in 4% paraformaldehyde (PF) in PBS for 25 min at room temperature. Adult midguts were dissected similarly and fixed for 40 min. Fixed tissues were washed three times in PBS and permeabilized in PBST. Samples were then blocked in 10% BSA for 2 hrs, followed by overnight incubation with primary antibodies at 4°C (**Table S5**). After four washes in PBST, tissues were incubated with fluorescent secondary antibodies (Invitrogen, **Table S5**, 1:250 dilution) at room temperature for 2 hrs, protected from light. Tissues were counterstained with TO-PRO-3, DAPI, and/or Phalloidin as appropriate, washed in PBST and PBS, and mounted in Vectashield. Larval tracheae were also counterstained with Calcoflour white for chitin staining (Flaven-Pouchon and Moussian 2022). For ASP cell visualization, specifically, wandering late third-instar larvae were selected for dissection (Guha et al. 2008). Confocal imaging was performed on a Leica SP5, Leica Stellaris, Zeiss LSM700, or Nikon AXR, selected based on availability and experimental requirements. Images were processed and analyzed using Leica LAS AF software or FIJI.

**Fluorescence Intensity Measurements and CTCF Calculation**

All image acquisitions, particularly those involving fluorescence intensity measurements, were performed under identical parameters for control and experimental samples. Fluorescence intensity was quantified using FIJI. The freehand tool was used to select the entire cell or the cell membrane boundary (depending on whether the protein of interest showed cytoplasmic or membrane localization) and to measure surface area and integrated density. Background fluorescence was measured from a cell-free region devoid of a specific signal. The corrected total cell fluorescence (CTCF) was calculated using the formula:

CTCF = Integrated Density − (Area of Selected Cell × Mean Background Fluorescence)

(<https://theolb.readthedocs.io/en/latest/imaging/measuring-cell-fluorescence-using-imagej.html>).

All images were assembled, and figures prepared using Adobe Photoshop; illustrations were created in BioRender.

**Visualization of Reporters**

To visualize the 10XSTAT92E-GFP (Bach et al. 2007), TRE-dsRed (Chatterjee and Bohmann 2012) and tGPH (Britton et al. 2002) reporters, *Drosophila* larvae and adult tissues were dissected in chilled 1X PBS, fixed in 4% PF/PBS, and washed three times in PBS. To visualize the LDH-LacZ reporter (Lavista-Llanos et al. 2002), larval trachea were dissected and processed similarly. LDH-LacZ expression was detected using an anti-β-galactosidase antibody as described in the immunofluorescence protocol above. After counterstaining with TO-PRO-3/DAPI for 30 minutes, samples were washed three times in PBS, mounted in Vectashield, and imaged.

**Assay for cellular stress**

*DHE (ROS) staining.* ROS in imaginal discs were detected using Dihydroethidium (DHE) as described previously (Owusu-Ansah et al. 2008). Larvae were dissected in Schneider's medium (SM), and DHE (30nM, reconstituted in DMSO) was added to SM. Samples were incubated for 5 min, washed thrice with SM, and fixed in 4% PF/PBS for 7 min. After a PBS rinse, imaginal discs were dissected and mounted in Vectashield. Samples were imaged immediately to avoid DHE oxidation.

*Glucose uptake assay using 2-NBDG.* Glucose uptake was assessed using the fluorescent glucose analog 2-NBDG (2-(N-(7-Nitrobenz-2-oxa-1,3-diazol-4-yl) Amino)-2-Deoxyglucose) (Zou et al. 2005). As described previously (de la Cova et al. 2014), imaginal discs obtained from late third instar larva (110 hr after egg laying) were carefully dissected in PBS and incubated in 2-NBDG (0.3mM in PBS) for 15 minutes in the dark at room temperature. Following three PBS washes, the samples were fixed in 4% PF for 15 min, mounted in an antifade medium, and imaged using confocal microscopy with standard FITC filter settings.

**DNA damage estimation**

*8-oxo-dG estimation.* The buildup of 8-oxo-2′-deoxyguanosine (8-oxo-dG), a well-established indicator of nucleotide oxidation, was detected as previously described (Kizhedathu et al. 2021). Late third instar larvae were dissected in ice-cold 1XPBS, fixed in 4% PF, and the imaginal disc of the desired genotype was immunostained as described (Kizhedathu et al. 2021)

*RPA70-GFP reporter.* We assessed the occurrence of single-stranded DNA breaks in wing imaginal discs using a fly strain that constitutively expresses the RPA70‑GFP fusion protein (Blythe and Wieschaus 2015).  Under normal conditions, RPA70 is evenly distributed throughout the nucleus, but upon single-strand DNA damage, it localizes to discrete nuclear foci that mark the DNA break sites (Blythe and Wieschaus 2015). For the RPA70-GFP assessment, staining was done as described above.

**TUNEL assay for cell death**

The third instar larval wing discs were dissected as previously described (Dekanty et al. 2012) and fixed in 4% PF. The TUNEL assay was performed using the Click-iT™ Plus TUNEL Assay (Invitrogen) according to the manufacturer's protocol.

**Drug feeding in adult flies**

The Akt inhibitor (GSK690693) (Cheng et al. 2022) was fed to adult flies to assess rescue of cut-wing phenotype. Briefly, the 5µM and 15µM concentrations of the drug was mixed with standard fly food (Markstein et al. 2014). For control, DMSO was added to the fly food. 25 adult flies per vial were allowed to feed on the drug-containing food, and the vials were changed every 3 days.

**Smurf assay for intestinal epithelial barrier**

For the Smurf assay, an intestinal barrier defect assay (Rera et al. 2012), flies were reared on regular food at room temperature for five days. Flies were fed blue food (2.5% bromophenol blue) for 12 hrs and directly screened for the Smurf phenotype when the dye coloration was seen outside the digestive tract. Images were acquired using a Leica M205 FA stereomicroscope.

*Gut Cell Area and Perimeter*. Segmentation analysis was performed using PYTHON as described below to identify cell area and perimeter. The resulting pixel-based measurements were exported in CSV files and plotted in GraphPad Prism 8.0 and analyzed by paired t-test.

**Tracheal cell size quantification and statistical analysis**
To calculate individual cell areas of flattened tracheal cells around a tubular luminal structure, we separately measured the cell areas from the upper half of the projection (top to the middle of the lumen) and the remaining lower half of individual coracle-lined cell boundaries using ImageJ, which were then added to derive their final cell sizes. Cell area and fluorescence intensity were calculated using Image J (FIJI). A freehand drawing tool was selected to trace the cell or nuclear boundaries and calculate its surface area and integrated density. For junctional length, the line tool was used to trace the width of the junction marked by the coracle.

**3D visualization and Morphometric Analysis**
Three-dimensional rendering and lumen quantification were performed using ImageJ (FIJI). Confocal Z-stacks were acquired to encompass the entire tracheal diameter, capturing the epithelium and the central lumen. To visualize the architecture, 3D projections were generated using the "Stacks" tool, with rotations along the X or Y axis relative to the tracheal orientation. For volumetric visualization, 3D videos were produced via the "3D Viewer" plugin.

**Mathematical modeling of the impact of squamous-to-cuboidal transition on tracheal tube geometry**

*Assumptions.* We model the tracheal wall as an elastic epithelial sheet folded into a cylinder. We assume cells can freely rearrange in-plane, meaning the sheet offers no in-plane shear resistance but resists out-of-plane bending. Because the tube is tethered to anatomical landmarks, it has a preferred axial length (*L_0_*), where any deviation requires mechanical work. The final radius (*R*) and length (*L*) are not fixed but emerge from an energetic balance between these constraints.

*Model Formulation.* Since the sheet is neither torn nor overlapped, its total area is considered to be conserved. When formed into a cylinder of radius $R$ and length $L$, the lateral surface area is

$$A=2\pi RL$$

This provides a useful constraint, allowing us to express the length as

$$L=\frac{A}{2\pi R}$$

Thus, the problem effectively reduces to determining the optimal radius $R$.

The epithelial sheet resists bending, and the associated energy depends on the curvature. For a cylinder, the bending energy per unit area for a thin elastic sheet is

$$u=\frac{1}{2}D\kappa^{2}$$

where $\kappa=1/R$ is the curvature and the bending rigidity $D$

$$D=\frac{Et^{3}}{12(1-\nu^{2})}$$

with $E$ being Young's modulus and $\nu$ the Poisson ratio of the epithelial cells.

In addition to bending, we introduce a preferred length $L_{0}$ for the tube, such that deviations from this length are penalized. This is modeled through a quadratic energy term:

$$U_{\text{length}}=\frac{1}{2}K(L-L_{0})^{2}=\frac{1}{2}K\left( \frac{A}{2\pi R}-L_{0} \right)^{2}$$

where $K$ is an effective stiffness associated with axial deformation. Therefore, the total energy of the system is

$$U(R)=\frac{1}{2}D\frac{A}{R^{2}}+\frac{1}{2}K\left( \frac{A}{2\pi R}-L_{0} \right)^{2}$$

To find the equilibrium configuration, we minimize this energy with respect to $R$. This yields the condition

$$\frac{D}{R^{3}}+\frac{KA}{4\pi^{2}R^{3}}-\frac{KL_{0}}{2\pi R^{2}}=0$$

which can be solved to obtain the equilibrium radius

$$R=\frac{2\pi D}{KL_{0}}+\frac{A}{2\pi L_{0}}=\frac{2\pi}{KL_{0}}\left( \frac{Et^{3}}{12(1-\nu^{2})} \right)+\frac{A}{2\pi L_{0}}$$

This result highlights a balance between two competing effects. The bending energy penalizes curvature and therefore favors larger radii, while the axial constraint enforces a preferred length, indirectly selecting a corresponding radius through the geometric constraint. Because the total area is fixed, increasing the radius necessarily shortens the cylinder, and deviations from the preferred length are energetically costly. The system settles into a configuration where these competing effects balance each other.

This framework illustrates the geometric consequences of a squamous-to-cuboidal transition. This transition increases cell height, and thus the effective thickness (*t*) of the epithelial sheet. Because bending rigidity scales cubically with thickness ( $D\sim t^{3}$), even modest height changes substantially increase bending resistance. To minimize this higher energetic penalty for curvature, the system shifts toward a larger radius. Given that total surface area (*A*) is conserved, the model predicts that this transition necessarily results in a shorter, wider tracheal tube (Landau et al. 1986)**.**

*Computer simulation (Vertex Model).* For the computer simulation of the tracheal epithelium, we consider it as a two-dimensional sheet discretized into a network of polygonal cells (similar to the vertex model (Farhadifar et al. 2007)), embedded in three-dimensional space and topologically arranged on a cylindrical surface. The configuration is specified by the positions of vertices $\boldsymbol{r}_{i}=\left( R,\theta_{i},z_{i} \right)$, which define cell shapes, edge connectivity, and surface geometry. Please note that the polar $\theta_{i}$ and axial $z_{i}$ components of the coordinate for each vertex are different, but all vertices share the radial $R$ component, which is the same as the cylinder radius. The total surface is periodic in the circumferential direction, forming a closed cylindrical topology, while remaining free along the axial direction

*Mechanics.* The in-plane mechanics are described by a standard vertex model energy

$$\begin{aligned} U_{\text{VM}}=\sum_{c} \left[ \frac{K_{A}}{2}\left( A_{c}-A_{c}^{0} \right)^{2}+\frac{K_{P}}{2}\left( P_{c}-P_{c}^{0} \right)^{2} \right] \end{aligned}$$

where $A_{c}^{0}$ and $P_{c}^{0}$ are preferred cell area and perimeter, and $K_{A}$, $K_{P}$ are elastic moduli. This formulation captures effective cell incompressibility and cortical tension.

Following the continuum description, the epithelial sheet resists curvature. For a cylindrical surface, the curvature is $\kappa=1/R$ and the bending energy is:

$$\begin{aligned} U_{\text{bend}}=\frac{1}{2}D\int\kappa^{2}dA. \end{aligned}$$

The total energy of the system trachea is, therefore, given by

$$\begin{aligned} U=U_{\text{VM}}+U_{\text{bend}}+U_{\text{axial}}+U_{\text{area}} \end{aligned}$$

where $U_{\text{axial}}$ and $U_{\text{area}}$ arise from the axial deformation and surface area change as described in the continuum model. We minimize this energy with the constraint of maintaining the cylindrical shape to obtain the equilibrium configuration of the cells.

The system evolves via overdamped dynamics

$$\begin{aligned} \gamma{\dot{\boldsymbol{r}}}_{i}=-\frac{\partial U}{\partial\boldsymbol{r}_{i}}, \end{aligned}$$

where $\gamma$ is a friction coefficient. Vertex positions are updated iteratively until convergence. Topological re-arrangements (T1 transitions) are implemented when edge lengths fall below a threshold, ensuring fluid-like in-plane behavior.

The vertex model captures cell-level mechanical properties such as area elasticity and cortical tension, while the bending term encodes resistance to curvature. Increasing epithelial thickness enhances bending rigidity ($D\sim t^{3}$), leading to reduced curvature and thus larger equilibrium radii, consistent with squamous-to-cuboidal transitions.

The detailed code for analysis of the trachea model can be found here <https://github.com/suhailrbme/tracheaAnalysisModel.git>

**Burrowing and Tunneling assay**
We performed the burrowing and tunnelling assay for primary screening of hypoxia induced by SARS-CoV-2 NSP3 or PLpro using the established protocol (Qiang et al. 2018; Zhou et al. 2016). *Drosophila* larvae burrow into the food with their posterior spiracles exposed at the surface of the food for gaseous exchange. The foraging behavior of larvae occurs in the third instar stage, where they dig into the food, form tunnels, and wander as pre-pupation activities. This larval movement is energetically costly; hence, hypoxic larvae do not burrow to save their energy reserves.

**Stable transfection of Madin Darby Canine Kidney (MDCK) by SARS-CoV-2 NSP3 PLpro**
NSP3 PLpro was codon-optimised and tagged with Myc at the C-terminus. The desired gene of interest, PLpro-Myc, was then ligated into the pLVX-TetOne-Puro vector (Takara Bio) using BamHI/EcoRI restriction enzymes and subsequently transformed into *E. coli* competent cells. Positive clones were identified, and the plasmid was isolated and sequenced for further confirmation. The pLVX-TetOne-Puro vector contains a Tet-On inducible expression system, in which the gene of interest is under tight control of the TRE3G promoter (PTRE3GS), and cells express the Tet-On 3G transactivator protein. The target cells thus express high levels of gene of interest when cultured in the presence of doxycycline (Dox). Stable transgenic MDCK cells were generated by Aurigene Pharmaceutical Services (Hyderabad, India) by lentiviral transduction as per the manufacturer's protocol. The MDCK cells were grown in a 37°C, 5% CO_2_ humid incubator using Minimal Essential Medium (MEM) containing Earl's salts and supplemented with 2mM L-glutamine, 1mM sodium pyruvate, 10% Fetal Bovine Serum, and Penicillin-Streptomycin or Anti-Anti.

**Immunohistochemistry, imaging, quantification, and statistical analysis**

Cells were seeded onto coverslips in 6-well plates with selection media (MEM, 10% FBS, 0.5 µg/ml puromycin) and incubated at 37°C and 5% CO₂. After 24 hrs, test wells were induced with 0.5 µg/ml doxycycline. Media containing doxycycline was replenished at 48 hr and cells were incubated for a further 24 hrs to maintain a continuous doxycyline induction for 72 hrs. Cells were washed with PBS, fixed with 4% PF for 10 min, rinsed, then blocked (4% BSA in 0.2% PBT) for 2 hrs. After washing with washing buffer (WB; 0.05% PBT), cells were incubated overnight with the primary antibody. The cells were washed and incubated with the secondary antibody for 1.5 hr at room temperature, followed by three washes with WB and two with PBS. Cells were counterstained with TO-PRO-3 for 5 minutes at RT in the dark to mark the nucleus. Cells were mounted in Vectashield® and imaged in a confocal microscope with a 63X oil-immersion objective. Images were analyzed and processed using Leica confocal software, LAS AF, or FIJI.

3D images of E-cadherin were generated in FIJI using 3D projection of z-stacks.

CTCF was calculated as previously described. For H2AX and LC3B puncta calculation, the image threshold was set to a constant value for all images. From the Analyze particle dropdown, the size of the particle and circularity were set to avoid background puncta. All statistical analyses were performed in GraphPad 8.0. Unpaired *t*-tests were performed to quantify fluorescence intensities and the number of puncta. A cutoff of *p*<0.05 was used to define statistical significance in all graphical plots. Illustrations were made using BioRender.

*Segmentation analysis.* Confocal images of MDCK monolayers were analyzed using maximum intensity projections of *z*-stack images. Cell segmentation and area quantification were performed using custom Python scripts. The cells were stained with ZO-1 to mark the cell boundaries. The ZO-1 signal was used to set the threshold, and the boundary skeletons were generated, ignoring the outermost cells. Cells were colored with the jet colormap to distinguish cell sizes. The resulting pixel-based measurements were exported in CSV files and plotted in GraphPad Prism 8.0 and analyzed by paired t-test. The detailed code can be found here <https://github.com/suhailrbme/tracheaAnalysisModel.git>

*Junctional Intensity Measurement.* The intensity of junctional proteins across the apical plane of the junction (horizontal measurement) was plotted using ggplot2 in RStudio, as described earlier (Lovejoy et al. 2025). For horizontal TJ/AJ quantification, Z-stacks were projected using a sum-intensity projection of all relevant planes into a composite image. ROIs were then drawn perpendicular to the junctional line in Fiji/ImageJ. For each ROI, the ImageJ macro generated fluorescence profiles for the TJ marker (e.g., ZO-1) and the AJ marker (e.g., E-cadherin), producing one file for each ROI and channel for downstream analysis in R. The code for the ImageJ macro and RStudio was used as mentioned previously (Lovejoy et al. 2025). The significance test was performed using one-way ANOVA.

*DHE (ROS), and Lysotracker staining.* Growth media was removed from the cells after 72 hrs of doxycycline induction, and the cells were washed with PBS. Live cells were stained with DHE (5 µM) (Kumar and Gullapalli 2024), or Lysotracker (75nM) (Huang et al. 2024) and incubated for 30 minutes at room temperature in the dark. Cells were washed with PBS, mounted with Antifade, and imaged immediately.

*TUNEL assay.* Growth media was removed from the cells after 72 hrs of doxycycline induction, and the cells were washed with PBS. The TUNEL assay was performed using the Click-iT™ Plus TUNEL Assay according to the manufacturer's protocol.

**Quantification of inflammatory cytokine expression in SARS-CoV-2 *PLpro*-expressing *Drosophila* tissue and MDCK cells**

Transcriptional upregulations of inflammatory cytokines upon SARS-CoV-2 *PLpro* expression in *Drosophila* larval wing imaginal discs, adult gut tissues, and MDCK cells were measured by quantitative real-time PCR (qPCR). For wing disc analysis, wild-type and *vg>PLpro* discs were dissected, while for gut studies, wild-type (*MyoIA^ts^>+* or *mex^ts^>+*) and *MyoIA^ts^>PLpro* or *mex^ts^>PLpro* transgenic guts were used. For MDCK cells, both doxycycline-induced and non-induced cells were used 72 hrs after doxycycline treatment. Total RNA was extracted from each group using the TRIzol reagent as described previously (Rio et al. 2010), and RNA concentrations were determined by UV spectrophotometry. Following standard protocols, 50 ng of RNA from each sample was reverse-transcribed into cDNA. qPCR was performed using SYBR Green chemistry on a Thermo Scientific QuantStudio 5 Real-Time PCR Instrument. Expression of cytokine genes (**Table S2**) was measured, with Actin as an internal control. Relative gene expression levels were calculated using the 2^−ΔΔCt^ method, where ΔCt values represent the difference between the target gene and Actin Ct values for each sample. For each genotype or condition, ΔCt values are the mean of three independent biological replicates ± 1 standard deviation. Fold changes in transcript levels were determined by comparing control samples to *PLpro*-expressing samples (Bajpai et al. 2020; Capellini et al. 2020).

**Table S1. Primers for sub-cloning the PLpro domain from NSP3.**

| PLPro | **Forward**: (NotI) | GAAGATCTATGGGCGAGGTGCG |
| --- | --- | --- |
|  | **Reverse**: (XhoI) | CGCTCGAGTTAGGTGGTGGTATAGCT |

**Table S2. List of qPCR-Primers for analysis of cytokine levels in *Drosophila* tissues**

| **Gene** | **Primers** | |
| --- | --- | --- |
| *Actin* | **Forward** | ACGGAGCGTGGCTACAGC |
|  | **Reverse** | TCCTGATGTCACGCACGA |
| *upd1* | **Forward** | TTCCACGGCACAGTCAAG |
|  | **Reverse** | ACTCAGCACCAGCATCAC |
| *upd2* | **Forward** | TGGAGGTCAGCTGGGAATACC |
|  | **Reverse** | TGCAAAATGTCAGGGAGAAGTA |
| *upd3* | **Forward** | TCCAGAACAACTATGAGGGTGA |
|  | **Reverse** | TCCTGATTCTTTACCTTGCTCTT |
| *eiger* | **Forward** | CGTCCATTCTTGCCCAAAC |
|  | **Reverse** | AGCCCTGAGCCCTTAATTC |

**Table S3. List of qPCR-Primers for analysis of cytokine levels in MDCK cell lines:**

| ***Gene*** | ***Primer*** | | ***Source*** |
| --- | --- | --- | --- |
| ACTB | **Forward** | ACGGAGCGTGGCTACAGC | (Sauter et al. 2005) |
|  | **Reverse** | TCCTGATGTCACGCACGA |  |
| IL-6 | **Forward** | TCCAGAACAACTATGAGGGTGA | (Cavalcanti et al. 2015) |
|  | **Reverse** | TCCTGATTCTTTACCTTGCTCTT |  |
| STAT5 | **Forward** | TTGACTCTCCTGACCGCAAC | (Capellini et al. 2020) |
|  | **Reverse** | TCCGTCTACTGCTTTAGCGA |  |

**Table S4**. **Genetic stocks used**

| ***Stock*** | ***Source*** | ***Identifier*** |
| --- | --- | --- |
| *w^1118^* | BDSC | 5905 |
| *vg-Gal4* | Sean Carroll | (Kim et al. 1996) |
| *ptc-Gal4* | BDSC | 2017 |
| *en-Gal4* | S.C. Lakhotia | (Simmonds et al. 1995) |
| *MyoIA-Gal4; tub Gal80^ts^* | Sveta Chakrabarti | (Jiang et al. 2009) |
| *mex-Gal4; tub Gal80^ts^* | Sveta Chakrabarti | (Weaver et al. 2020) |
| *btl-Gal4* | Renjith Mathew | (Mathew et al. 2020) |
| *btl-DEcad-mtomato* | Marta Llimargas | (Casani et al. 2020) |
| *UAS GFP-nls* | BDSC | 4776 |
| *UAS-mRFP* | BDSC | 27392 |
| *UAS-SARS-CoV-2-nsp1* | BDSC | 94224 |
| *UAS-SARS-CoV-2-nsp2* | BDSC | 94243 |
| *UAS-SARS-CoV-2-nsp3* | BDSC | 94225 |
| *UAS-PLpro/TM6B* | This study | -- |
| *UAS-SARS-CoV-2-nsp4* | BDSC | 94245 |
| *UAS-SARS-CoV-2-nsp5* | BDSC | 95279 |
| *UAS-SARS-CoV-2-nsp6* | BDSC | 94247 |
| *UAS-SARS-CoV-2-nsp7* | BDSC | 94249 |
| *UAS-SARS-CoV-2-nsp8* | BDSC | 94250 |
| *UAS-SARS-CoV-2-nsp9* | BDSC | 94252 |
| *UAS-SARS-CoV-2-nsp10* | BDSC | 94232 |
| *UAS-SARS-CoV-2-nsp11* | BDSC | 94268 |
| *UAS-SARS-CoV-2-nsp12* | BDSC | 94234 |
| *UAS-SARS-CoV-2-nsp13* | BDSC | 94236 |
| *UAS-SARS-CoV-2-nsp14* | BDSC | 94238 |
| *UAS-SARS-CoV-2-nsp15* | BDSC | 94240 |
| *UAS-SARS-CoV-2-nsp16* | BDSC | 94242 |
| *UAS-SARS-CoV-2-ORF3a* | BDSC | 94226 |
| *UAS-SARS-CoV-2-ORF3b* | BDSC | 94256 |
| *UAS-SARS-CoV-2-ORF6* | BDSC | 94258 |
| *UAS-SARS-CoV-2-ORF7a* | BDSC | 94228 |
| *UAS-SARS-CoV-2-ORF7b* | BDSC | 94260 |
| *UAS-SARS-CoV-2-ORF8* | BDSC | 94262 |
| *UAS-SARS-CoV-2-ORF9b* | BDSC | 94264 |
| *UAS-SARS-CoV-2-ORF9c* | BDSC | 94266 |
| *UAS-SARS-CoV-2-ORF10* | BDSC | 94254 |
| *UAS-SARS-CoV-2-kozak nsp3* | BDSC | 98458 |
| *UAS-SARS-CoV-2-nsp3.C857A (mutant)* | BDSC | 98459 |
| *UAS-myrAKT* | Hugo Stocker | (Stocker et al. 2002) |
| *UAS Dronc^DN^* | BDSC | 58992 |
| *UAS-AKT RNAi* | BDSC | 33615, 82957 |
| *puc-GFP* | BDSC | 23694 |
| TRE-dsRed | BDSC | 59011 |
| 10XSTAT92E-GFP | BDSC | 26197 |
| *UAS Duox RNAi* | Arjun Guha | (Kizhedathu et al. 2021) |
| *RPA70-EGFP* | Arjun Guha | (Blythe and Wieschaus 2015) |
| *LDH LacZ* | Pablo Wappner | (Lavista-Llanos et al. 2002) |
| *UAS bsk-RNAi* | BDSC | 32977 |
| *UAS Stat-RNAi* | BDSC | 33637 |

**Table S5. KEY RESOURCES TABLE: Reagents and resources**

| ***Antibodies and stains*** | ***Source*** | ***Identifier*** |
| --- | --- | --- |
| Mouse Anti-Coracle | DSHB | C566.9 |
| Rabbit Anti-4EBP1 | Cell Signaling | 2855 |
| Mouse Anti-MMP1 | DSHB | 3A6B4 |
| Rabbit Anti-Snakeskin | Gift from Yasushi Izumi | (Izumi et al. 2012) |
| Mouse anti-β-Gal | Sigma-Aldrich | G6282 |
|  | DSHB | 40-1a |
| Rat Anti-DE-Cad | DSHB | DCAD2 |
| Mouse Anti-Myc tag | Cell Signaling Technology | 2276T |
| Rabbit Anti-ZO-1 | Invitrogen | 61-7300 |
| Mouse Anti-Occludin | Invitrogen | OC-3F10 |
| Rabbit Anti-Phospho-H2AX-S139 | Abclonal | AP0687 |
| Rabbit Anti-Caspase 3 | Sigma-Aldrich | C8487 (Fogarty and Bergmann 2014) |
| Mouse Anti-8-oxo-dG | Gift from Arjun Guha | Abclonal Cat #ab48508 |
| Rabbit Anti-E-Cad | Cell Signaling | 24E10 |
| Rabbit Anti PLpro | GeneTex | GTX638553 |
| Alexa Fluor 555 Goat Anti-Rabbit IgG | Invitrogen | A32732 |
| Alexa Fluor 633 Goat Anti-Rabbit IgG | Invitrogen | A31576 |
| Alexa Fluor 555 Goat Anti-Mouse IgG | Invitrogen | A32727 |
| Alexa Fluor 633 Goat Anti-Mouse IgG | Invitrogen | A21052 |
| Alexa Fluor 555 Goat Anti-Rat IgG | Invitrogen | A21434 |
| 2-NBDG (2-(N-(7-Nitrobenz-2-oxa-1,3-diazol-4-yl) Amino)-2-Deoxyglucose) | Sigma-Aldrich | N13195 |
| Schneider′s Insect Medium | HiMedia | IML003A |
| Click-iT™ Plus TUNEL Assay | Gift from Jonaki Sen | Invitrogen C10617 |
| Lysotracker | Invitrogen | DND-99 |
| Rhodamine Dextran | Invitrogen | D22912 |
| FD&C; BLUE No. 1 | Sigma-Aldrich | 861146 |
| TO-PRO-3 | Invitrogen | S33025 |
| DAPI | Invitrogen | S33025 |
| Calcoflour white stain | Sigma-Aldrich | 18909-100ML-F |
| Phalloidin-633 | Invitrogen | A22284 |
| Phalloidin-555 | Invitrogen | A34055 |
| Phalloidin-488 | Abcam | AB176753 |
| ProLong™ Gold | Invitrogen | P36930 |
| Vectashield | Vector Labs | H-1200 |
| Akt inhibitor | Sigma | GSK690693 |
| ***Software and Algorithms*** |  |  |
| Leica LAS AF software | Leica microsystem | http://www.leicamicrosystems.com/productes/microscope-software/ |
| FIJI | ImageJ | https://imagej.net/software/fiji/ |
| Prism | Graph pad | https://www.graphpad.com |
| Photoshop | Adobe CC | http://www.adobe.com |
| Illustrator | Adobe CC | http://www.adobe.com |
| BioRender | BioRender | https://biorender.com/ |
| RStudio R 3.6.0+ | RStudio 2026 | https://posit.co/download/rstudio-desktop/ |
| **Cell line Reagents** |  |  |
| MDCK cells | Aurigene |  |
| Minimum Essential Medium (1X) | Gibco | 11090-073 |
| FBS | Hyclone | SH30071.03 |
| Penicillin-Streptomycin | Invitrogen | 15140122 |
| Anti-Anti | Gibco | 15240062 |
| Puromycin | Sigma-Aldrich | P8833 |
| Doxycycline | Sigma-Aldrich | D3072 |
| **Other** |  |  |
| Leica TCS SP5 microscope | Leica microsystem | http://www.leica-microsystems.com/ |
| Leica Stellaris microscope | Leica microsystem | http://www.leica-microsystems.com/ |
| Leica M205 FA stereomicroscope | Leica microsystem | http://www.leica-microsystems.com/ |
| AXR Nikon Confocal Microscope | NIS elements | https://www.microscope.healthcare.nikon.com |
| Zeiss LSM700 Confocal Microscope | Zeiss | https://www.zeiss.com/corporate/en/products-and-solutions.html |

Landau, Lev Davidovich, L. D. Landau, Evgeniĭ Mikhaĭlovich Lifshit︠s︡, A. M. Kosevich, E. M. Lifshitz, and L. P. Pitaevskii. 1986. *Theory of Elasticity: Volume 7*. Elsevier.

Lavista-Llanos, Sofía, Lázaro Centanin, Maximiliano Irisarri, et al. 2002. “Control of the Hypoxic Response in Drosophila Melanogaster by the Basic Helix-Loop-Helix PAS Protein Similar.” *Molecular and Cellular Biology* 22 (19): 6842–53. https://doi.org/10.1128/MCB.22.19.6842-6853.2002.

Lovejoy, Madeline, Claudio A. Mucci, Agustín Rabino, and Rafael Garcia-Mata. 2025. “Protocol for Quantifying the Horizontal and Vertical Distribution of Junctional Proteins in Fixed Epithelial Cells.” *STAR Protocols* 6 (3): 104079. https://doi.org/10.1016/j.xpro.2025.104079.

Marchetti, Marco, Chenge Zhang, and Bruce A. Edgar. 2022. “An Improved Organ Explant Culture Method Reveals Stem Cell Lineage Dynamics in the Adult Drosophila Intestine.” *eLife* 11 (August): e76010. https://doi.org/10.7554/eLife.76010.

Markstein, Michele, Samantha Dettorre, Julio Cho, Ralph A. Neumüller, Sören Craig-Müller, and Norbert Perrimon. 2014. “Systematic Screen of Chemotherapeutics in Drosophila Stem Cell Tumors.” *Proceedings of the National Academy of Sciences of the United States of America* 111 (12): 4530–35. https://doi.org/10.1073/pnas.1401160111.

Mathew, Renjith, L. Daniel Rios-Barrera, Pedro Machado, Yannick Schwab, and Maria Leptin. 2020. “Transcytosis via the Late Endocytic Pathway as a Cell Morphogenetic Mechanism.” *The EMBO Journal* 39 (16): 16. https://doi.org/10.15252/embj.2020105332.

Morgan, Nancy Strom, Daniel M. Skovronsky, Spyridon Artavanis-Tsakonas, and Mark S. Mooseker. 1994. “The Molecular Cloning and Characterization of *Drosophila Melanogaster* Myosin-IA and Myosin-IB.” *Journal of Molecular Biology* 239 (3): 347–56. https://doi.org/10.1006/jmbi.1994.1376.

Owusu-Ansah, Edward, Amir Yavari, and Utpal Banerjee. 2008. “A Protocol for in Vivo Detection of Reactive Oxygen Species.” *Protocol Exchange*, ahead of print, February 27. https://doi.org/10.1038/nprot.2008.23.

Qiang, Karen M., Fanli Zhou, and Kathleen M. Beckingham. 2018. “A Burrowing/Tunneling Assay for Detection of Hypoxia in Drosophila Melanogaster Larvae.” *Journal of Visualized Experiments : JoVE*, no. 133 (March): 133. https://doi.org/10.3791/57131.

Rera, Michael, Rebecca I. Clark, and David W. Walker. 2012. “Intestinal Barrier Dysfunction Links Metabolic and Inflammatory Markers of Aging to Death in Drosophila.” *Proceedings of the National Academy of Sciences of the United States of America* 109 (52): 21528–33. https://doi.org/10.1073/pnas.1215849110.

Rio, Donald C., Manuel Ares, Gregory J. Hannon, and Timothy W. Nilsen. 2010. “Purification of RNA Using TRIzol (TRI Reagent).” *Cold Spring Harbor Protocols* 2010 (6): pdb.prot5439. https://doi.org/10.1101/pdb.prot5439.

Sauter, S. N., K. Allenspach, F. Gaschen, A. Gröne, E. Ontsouka, and J. W. Blum. 2005. “Cytokine Expression in an Ex Vivo Culture System of Duodenal Samples from Dogs with Chronic Enteropathies: Modulation by Probiotic Bacteria.” *Domestic Animal Endocrinology* 29 (4): 605–22. https://doi.org/10.1016/j.domaniend.2005.04.006.

Scholl, Aaron, Istri Ndoja, and Lan Jiang. 2021. “Drosophila Trachea as a Novel Model of COPD.” *International Journal of Molecular Sciences* 22 (23): 12730. https://doi.org/10.3390/ijms222312730.

Schulz, Robert A., Xiaoling Xie, Andrew J. Andres, and Samuel Galewsky. 1991. “Endoderm-Specific Expression of the *Drosophila Mex1* Gene.” *Developmental Biology* 143 (1): 206–11. https://doi.org/10.1016/0012-1606(91)90068-E.

Shiga, Yasuhiro, Miho Tanaka-Matakatsu, and Shigeo Hayashi. 1996. “A Nuclear GFP/β-Galactosidase Fusion Protein as a Marker for Morphogenesis in Living Drosophila.” *Development, Growth & Differentiation* 38 (1): 99–106. https://doi.org/10.1046/j.1440-169X.1996.00012.x.

Simmonds, Andrew J., William J. Brook, Stephen M. Cohen, and John B. Bell. 1995. “Distinguishable Functions for Engrailed and Invected in Anterior–Posterior Patterning in the Drosopila Wing.” *Nature* 376 (6539): 424–27. https://doi.org/10.1038/376424a0.

Simmonds, Andrew J., Xiaofeng Liu, Kelly H. Soanes, Henry M. Krause, Kenneth D. Irvine, and John B. Bell. 1998. “Molecular Interactions between Vestigial and Scalloped Promote Wing Formation in Drosophila.” *Genes & Development* 12 (24): 3815–20. https://doi.org/10.1101/gad.12.24.3815.

Stocker, Hugo, Mirjana Andjelkovic, Sean Oldham, et al. 2002. “Living with Lethal PIP3 Levels: Viability of Flies Lacking PTEN Restored by a PH Domain Mutation in Akt/PKB.” *Science* 295 (5562): 2088–91. https://doi.org/10.1126/science.1068094.

Weaver, Lesley N., Tianlu Ma, and Daniela Drummond-Barbosa. 2020. “Analysis of Gal4 Expression Patterns in Adult Drosophila Females.” *G3: Genes|Genomes|Genetics* 10 (11): 4147–58. https://doi.org/10.1534/g3.120.401676.

Zhou, Fanli, Karen M. Qiang, and Kathleen M. Beckingham. 2016. “Failure to Burrow and Tunnel Reveals Roles for Jim Lovell in the Growth and Endoreplication of the Drosophila Larval Tracheae.” *PLOS ONE* 11 (8): 8. https://doi.org/10.1371/journal.pone.0160233.

Zou, Chenhui, Yajie Wang, and Zhufang Shen. 2005. “2-NBDG as a Fluorescent Indicator for Direct Glucose Uptake Measurement.” *Journal of Biochemical and Biophysical Methods* 64 (3): 207–15. https://doi.org/10.1016/j.jbbm.2005.08.001.

**SUPPLEMENTARY FIGURE LEGENDS:**

**Supplementary Figure 1. Structural organization of the SARS-CoV-2 genome, polyprotein processing, and NSP3/PLpro domain architecture.**

**(A)** Schematic map of the SARS-CoV-2 viral genome ($5^{'}$ to $3^{'}$) and the polyproteins pp1a and pp1ab. The viral cysteine proteases PLpro (encoded within NSP3) and NSP5 cleave the polyproteins at their respective target recognition sequences ($LXGG\downarrow X$ and $X\text{-}L/I/V/F/M\text{-}Q\downarrow A/S/G$) to release individual functional units.

(**Aʹ)** Representative layout of the 16 nonstructural proteins (Nsps) and 9 accessory proteins utilized in the unbiased *Drosophila* genetic screen. The transgene collection was generated in the Bellen laboratory (Guichard et al., 2023).

**(B)** Domain organization of the full-length host-injury factor NSP3 (~212 kDa), highlighting the sequential arrangement of the N-terminal ubiquitin-like domain 1 (ubl1), macrodomain, SARS-unique domain (SUD), ubiquitin-like domain 2 (ubl2), papain-like protease domain (PLpro, ~35 kDa), nucleic acid-binding domain (NAB), and the C-terminal ectodomain.

**(B')** Detailed structural schematic of the isolated PLpro catalytic domain fragment (residues 746–1060 of NSP3). The fragment consists of the N-terminal ubl2 domain (purple) followed by the Thumb (orange), Fingers (blue), and Palm (green) subdomains. The catalytic center is maintained by the conserved triad consisting of $C111$, $H272$, and $D286$, which executes viral polyprotein proteolysis, deubiquitination (DUB), and deISGylation.

**Supplementary Figure 2. Genetic screen identifies SARS-CoV-2 PLpro as a potent inducer of epithelial defects in the *Drosophila* wing.**

**(A)** Structural schematic of the *Drosophila* third-instar wing imaginal disc (left) and adult wing (right). Colors outline the distinct targeted driver domains: *vestigial-Gal4* (*vg-Gal4*; red) marking the prospective adult wing margin and *patched-Gal4* (*ptc-Gal4*; green) marking a narrow stripe running along the anterior–posterior (A–P) compartment boundary.

**(B)** Wing imaginal discs expressing nuclear GFP driven by *vg-Gal4* (left) or *ptc-Gal4* (right) to validate spatial tracking parameters.

**(C)** Adult wing phenotypes from *vg>GFP* (control) and *vg>reaper* (*vg>rpr*) flies. Dashed red lines trace the classic cut-wing margin degradation phenotype resulting from *reaper* expression, serving as a positive control for cell death induction.

**(D)** *ptc>GFP, PLpro* wing imaginal disc showing a merged composite (left) and the isolated red channel (right) immunostained with an anti-PLpro antibody to confirm driver-restricted expression.

**(E)** Adult wings following *vg-Gal4*-mediated expression of individual SARS-CoV-2 nonstructural proteins (NSPs) and accessory factors (ORFs) from the comprehensive transgenic library. Of the 25 screens, only NSP3 and NSP5 elicit the characteristic cut-wing margins (as tracked by dashed red lines).

**(F)** Quantitative box plot showing the percentage loss of total adult wing surface area across the key experimental groups (*Control*, *vg>NSP3*, and *vg>NSP5*).

**(G)** Wing imaginal discs from *vg>GFP* control (top) and *vg>NSP3; GFP* (bottom) larvae immunostained for active cell death using an anti-cleaved Caspase-3 antibody (red; labeled as Cas-3).

Data are presented as mean ± SEM. Statistical significance in **(F)** was evaluated via a one-way ANOVA followed by Tukey's multiple-comparison post-hoc test (****P<0.001*). Scale bars: 50 μm **(B, D)**, 25 μm **(G)**, and 0.5 mm **(C, E)**. Nuclei are counterstained with DAPI (blue) in **(B, D, G)**. For tissue arrays, n = 15 wing discs per genotype **(B, D, G)**, and n = 40 adult wings per genotype **(C, E)**.

**Supplementary Figure 3. PLpro-induced epithelial injury triggers a non-autonomous compensatory proliferative response.**

**(A)** Representative adult wings from *ptc>GFP* (control) and *ptc>GFP, PLpro* flies. Insets show high-magnification views of the regions outlined by red boxes. PLpro expression results in the complete loss of the anterior cross-vein (indicated by the red asterisk) and a reduction in the intervein space between longitudinal veins L2 and L3 (red line) compared to the intact control features (blue asterisk and blue line).

**(B)** Wing imaginal discs from *ptc>GFP* (control) and *ptc>GFP,PLpro* larvae immunostained for phospho-histone H3 (PH3; red), a marker of mitotic cells.

**(C)** Quantification of the total number of PH3-positive cells within the cell-autonomous driver domain ($\text{ptc}^{+}$) and the adjacent, non-transgene-expressing host tissue ($\text{ptc}^{-}$). PLpro expression induces non-autonomous over-proliferation in surrounding wild-type cells ($\text{ptc}^{-}$) despite a cell-autonomous reduction in mitosis within the injury zone ($\text{ptc}^{+}$).

Data are presented as mean ± SEM. Statistical significance was evaluated using a two-tailed Student's $t$-test (**P<0.05*; *****P<0.001*. Scale bars: 0.5 mm **(A)** and 25 μm **(B)**. For sample sizing, n=15 wing discs per genotype for **(B)**, and n = 40 adult wings per genotype for **(A)**. DAPI (blue) counterstains nuclei in **(B)**.

**Supplementary Figure 4. Genetic and pharmacological modulation of host Akt signaling phenocopies and mitigates PLpro-induced epithelial damage.**

**(A)** Wing imaginal disc expressing constitutively active myristoylated Akt (*ptc>GFP, myrAkt*) showing a merged composite (left) and the isolated red channel (right) immunostained for cleaved Caspase-3 (red; labeled as Cas-3).

**(B)** Overview (left) and high-magnification inset (right) of a representative adult wing from a *ptc>GFP, myrAkt* fly display reduction in the intervein region between longitudinal veins L2 and L3 (red line), along with complete structural loss of the anterior cross-vein between L3 and L4 (indicated by the red asterisk), phenocopying the architectural defects induced by PLpro.

**(C)** Adult wings from *ptc>GFP, PLpro* flies following developmental treatment with either a vehicle control (DMSO) or varying concentrations (5 µM and 15µM) of the specific small-molecule host Akt inhibitor GSK690693.

**(D)** Quantitative box plot detailing the percentage of adult flies exhibiting the characteristic cut-wing margin degradation phenotype across the indicated experimental treatment arms. Pharmacological inhibition of Akt signaling protects epithelial integrity and suppresses wing pattern loss in a dose-dependent manner.

Scale bars: 25 μm **(A)** and 0.5 mm **(B, C)**. DAPI (blue) counterstains nuclei in **(A)**. Statistical significance in **(D)** was evaluated via a one-way ANOVA followed by Tukey's multiple-comparison post-hoc test (**P<0.05*; *****P<0.0001*. For sample sizing, n = 15 wing imaginal discs per genotype for **(A)**, n = 40 adult wings per genotype for **(B, C)**, and n = 60 adult flies per group for **(D)**.

**Supplementary Figure 5.** **The gut epithelium of *Drosophila* adults and PLpro-induced gut dysfunction.**

**(A)** Quantification of Smurf phenotype in *MyoIA^ts^>+* (control), *mex^ts^>+* (control), *MyoIA^ts^>NSP3*, *MyoIA^ts^>PLpro,* and *mex^ts^>PLpro* adult flies’ incidence across indicated genotypes.

**(B)** Representative images of adult SMURF flies for the above genotypes.

Data are presented as mean ± SEM. Statistical significance in (A) was determined using a one-way ANOVA followed by Tukey's multiple-comparison post-hoc test (*P<0.05*; ****P<0.001*. Scale bars: 1 mm (B). For sample sizing, $n=40$ adult flies per genotype.

**Supplementary Figure 6. PLpro activates JAK–STAT and JNK signaling and promotes enterocyte hypertrophy in the adult midgut.**

**(A–C)** Posterior midgut epithelium from 10-day-old *mex^ts^>+* (control) and *mex^ts^>PLpro* adults stained for Snakeskin (Ssk; gray) (**A**). Quantification of enterocyte cell area (**B**) and cell perimeter (**C**).

**(D–F)** Posterior midgut epithelium from 10-day-old *MyoIA^ts^>+* (control) and *MyoIA^ts^>PLpro* adults stained for Snakeskin (Ssk; gray) (**D**). Quantification of enterocyte cell area (**E**) and cell perimeter (**F**).

**(G)** RT–qPCR analysis of *upd1*, *upd2*, *upd3*, and *eiger* transcript levels in midguts of 10-day-old *MyoIA^ts^>PLpro* adults relative to age-matched controls.

**(H–H′)** Adult midguts from *MyoIA^ts^>+* (control) and *MyoIA^ts^>PLpro* adults carrying the 10XSTAT92E-GFP reporter (green), reporter of JAK–STAT signaling. Quantification of 10XSTAT92E-GFP fluorescence intensity (**H′**).

**(I–I′)** Adult midguts from *MyoIA^ts^>+* (control) and *MyoIA^ts^>PLpro* adults carrying the TRE-dsRed reporter (red), reporter of JNK signaling. Quantification of TRE-dsRed fluorescence intensity (**I′**).

**(J)** Posterior midgut epithelium from 10-day-old *MyoIA^ts^>PLpro, STAT-RNAi* adults stained for Snakeskin (Ssk; gray). Smurf assay of *MyoIA^ts^>PLpro, STAT-RNAi* adults.

Data are presented as mean ± SEM. Scale bars: 20 μm (A, D, H, I, J). DAPI (blue) marks nuclei in (H, I). n = 15 midguts per genotype for (A, D, H, I, J) and n = 60 adults per genotype for the Smurf assay.

**Supplementary Figure 7. PLpro induces oxidative stress and JNK signaling in the adult midgut.**

**(A-B)** Posterior midgut epithelium from 10-day-old *mex^ts^>+* (control) and *mex^ts^>PLpro* adults stained with dihydroethidium (DHE; red) (**A**) and MMP1 (magenta) (**B**).

Scale bars: 20 μm. n = 15 midguts per genotype.

**Supplementary Figure 8. Characterization of the larval tracheal system and PLpro expression in tracheal epithelial cells.**

**(A)** Schematic representation of the third-instar larval tracheal system. The sixth dorsal trunk metamere (Tr6), highlighted in the diagram, was used for analyses of tracheal epithelial architecture throughout this study (adapted from (Scholl et al. 2021)).

**(A′)** Cross-sectional schematic of the dorsal trunk showing the tracheal lumen, taenidia (cuticular ridges that reinforce the lumen), and the surrounding squamous epithelial cells (adapted from (Harrison 2009)).

**(B)** Third-instar larva expressing *btl>GFP*, showing trachea-specific GFP expression.

**(C)** Third-instar larval dorsal tracheal trunk (DT) expressing DE-cadherin (gray), showing tracheal metameres Tr1–Tr8. Tr2 was used for analyses of tracheoblast biology, whereas Tr6 was used for assessment of tracheal epithelial architecture.

**(D)** Dorsal tracheal trunk (DT) cells from *btl>+* (control) and *btl>PLpro* larvae immunostained with an anti-PLpro antibody.

Scale bars: 1 mm (B), 200 μm (C), and 20 μm (D). n = 15 larvae per genotype.

**Supplementary Figure 9. Mathematical modeling of the PLpro-induced squamous-to-cuboidal transition in the larval trachea.**

**(A)** Vertex model simulations of tracheal tubes lined by squamous (top) or cuboidal (bottom) epithelial cells, generated using geometric parameters derived from experimental measurements of control and PLpro-expressing tracheae. Each model consists of 24 epithelial cells. The cuboidal cell-lined tube is shorter than the squamous cell-lined tube despite containing the same number of cells.

**(B)** Cross-sectional views of the simulated tracheal tubes shown in (A). The cuboidal cell-lined tube (right) exhibits an increased luminal perimeter and a thicker epithelial lining than the squamous cell-lined tube (left).

**Supplementary Figure 10. PLpro expression in the larval trachea induces hypoxic metabolic stress and JNK signaling.**

**(A)** Dorsal tracheal trunks (DTs) from third-instar *btl>+* (control) and *btl>PLpro* larvae carrying the LDH-lacZ reporter. β-galactosidase (β-Gal, green) indicates LDH reporter activity.

**(B)** Dorsal tracheal trunks (DTs) from third-instar *btl>+* (control) and *btl>PLpro* larvae carrying TRE-dsRed reporter (red).

Scale bars: 20 μm. *n* = 15 tracheae per genotype for (A, B).

**Supplementary Figure 11. PLpro induces oxidative stress, DNA damage, apoptosis, and inflammatory signaling in MDCK cells.**

**(A)** Immunoblot confirming PLpro (anti-Myc) expression in MDCK cells following doxycycline induction. β-actin serves as a loading control.

**(B)** Immunofluorescence of PLpro (anti-PLpro, red) in control and PLpro-expressing MDCK cells. Occludin (green) marks tight junctions.

**(C–C′)** Dihydroethidium (DHE; red) staining in control and PLpro-expressing MDCK cells (**C**). Quantification of DHE fluorescence intensity (**C′**).

**(D–D′)** Immunofluorescence of H2AX (green) and Occludin (red) in control and PLpro-expressing MDCK cells (**D**). Quantification of H2AX foci per nucleus (**D′**).

**(E–E′)** TUNEL staining (red) in control and PLpro-expressing MDCK cells (**E**). Quantification of TUNEL-positive cells (**E′**).

**(F)** RT–qPCR analysis of IL-6 and STAT5 transcript levels in PLpro-expressing MDCK cells relative to control.

Data are presented as mean ± SEM. ****P < 0.0001; **P < 0.01. Scale bars: 20 μm (B–E). DAPI (blue) marks nuclei in (D), and TO-PRO-3 (green) marks nuclei in (E). *n* = 5 independent experiments.

**Supplementary Video 1: Control larval tracheal architecture.** 3D reconstruction of a confocal Z-stack showing the dorsal tracheal trunk (segment Tr6) of a third-instar wild-type control larva (*btl-Gal4>+*). Septate junctions are marked by Coracle immunostaining, displaying typical elongated, flattened squamous cell boundaries with thin, uniform junctional bands.

**Supplementary Video 2: Tracheal remodeling induced by SARS-CoV-2 NSP3.** 3D reconstruction of a confocal Z-stack showing a *btl>NSP3* third-instar larval tracheal trunk (segment Tr6). Expression of full-length NSP3 drives an aberrant squamous-to-cuboidal cellular transition, characterized by a conspicuous reduction in apical cell area, nuclear enlargement, and dense thickening of Coracle-marked septate junctions.

**Supplementary Video 3: Tracheal remodeling induced by the isolated SARS-CoV-2 PLpro domain.** 3D reconstruction of a confocal Z-stack showing a *btl>PLpro* third-instar larval tracheal trunk (segment Tr6). Directed expression of the isolated protease domain alone phenocopies and exacerbates the full-length protein phenotype, leading to severe cell-intrinsic geometric packing defects and dense junctional accumulation of Coracle.

**Supplementary Video 4: Suppression of viral-induced tracheal defects by host JAK-STAT knockdown.** 3D reconstruction of a confocal Z-stack showing a *btl>NSP3, STAT-RNAi* third-instar larval tracheal trunk (segment Tr6). Knockdown of host JAK-STAT signaling restores the flattened squamous cell architecture and resets Coracle distribution to narrow, normal junctional bands.
